# Measurement and self-operating computer of the leukocyte continuum as a fixed space–time continuum in inflammation

**DOI:** 10.1101/543611

**Authors:** Yuusuke Nonomura

**Author notes:** To whom correspondence should be addressed: Yuusuke Nonomura, Nonomura Dental Clinic, National Yagata Bldg 2F 3-6, Higasiyamatoori, Chikusa-ku, Nagoya City, Aichi 4640807, Japan.

## Abstract

**Motivation:** No biomarkers and systems, including leukocyte count and flow cytometry, can be used to measure tissue injury for diagnosing inflammation. A fixed space–time continuum (SτC) biomarker can address this issue. A leukocyte continuum (LC) is a biomarker forming a SτC capable of measuring injury by operators and equations for a self-operating computation.

**Results:** A self-operating computer (SOC) LC as a water treatment for leukocyte(s) was generated using leukocyte(s). String leukocyte continuum (StrLC), single-layer leukocyte (SLL) and multilayer leukocyte continuum (MLC) were demonstrated in various LCs using an equation with a primitive-operator. In the SOC, the LC is the inflammation graph of the operation result. The relative differential equation (RDE) shows how to recognize the LC not as a ‘model’ in the conventional-other-operating-computer (cOOC), but as an actual arithmetic unit with a display unit. The SOC shows the essential nature in real time.

## INTRODUCTION

An other-operating computer (OOC) is a computer comprising NAND and NOR silicon logic chip that performs processing using algorithms on these circuits and displays graphs upon analogue-to-digital (A/D) conversion of sample information. Conventional natural phenomena are mainly analysed via cOOC, which considers the solution of the conventional convergence-differential equation (CDE) as the core (Example S1). However, the cOOC cannot calculate an absolute origin (AO) (singularity) at the boundary of multiple quadrants {an example is the problem of inconsistency in the directional properties associated with the equation-of-motion (IDP-EOM)}, different-object interference (AO) (singularity) (an example is the prey and predator problem) (Britton, 2010; Hawkins, 1999; Krainov, 2002; Teramoto, 2009; Thornley, 2007) and relative origin (RO) (singularity) (Saw, E.-W., et al., 2016). AO and RO do not have smoothness (Text S1). Their singularities are the origin of the problems that follow (e.g. non-calculable inflammation). Therefore, the conventional scientific method (e.g. Text S2-4, Material S1) only has the OOC until the end of this paper.

However, a self-operating computer (SOC) is the phenomenon itself that is recognized to be arithmetic circuit {by relative-differential equation: RDE (Definition S6)} with graphic display function {D() or D(N())} unlike the OOC. In SOC RDE and OOC by SOC, AO and RO (Definition S6.4) are calculable and smooth. {D() is an image device function (microscope, MRI, CT, telescope, etc.); N() is the function indicating diffusion extent from tissue injury and that of the samples (on glass slide) covered with a glass cover.

One example is the Navier–Stokes equation (NSE) generated using the RDE. The observation image is D(RDE) or D(N(RDE)). When sampling a tissue injury and making a direct observation, one directly observes the RDE arithmetic result. The SOC consists of the formation of a fixed space–time continuum (SτC) by an observed object and from the RDE that shows the SOC. The solution takes kOOC and zOOC. The SOC shows inflammation, elementary particle, life, and new figure of the space–time continuum. Inflammation is one such example of the SOC. The directional property that measures decomposed single cell, including cell injury, takes the direction of modern pathology, and is indicated in a (blood) cell-counting system (Graham et al., 2003), a cell measurement system as flow cytometry (FCM) (Cossarizza et al., 2017) and leukocyte counts (Lcnt) (Saito, 2002; DACIE, 1975). This is defined as “cell (injury) element theory” (CET). The method describes the royal road of the OOC consisting of decomposition into element (cell) from a sample and analysis (operation processing) of the sample. Per cell decomposition destroys the tissue structure (tissue injury). Cell injury cannot measure inflammation, which is the best biomarker of an invasive disease. However, it can be measured with tissue injury and understood by observations of tissue sections (Rubin, 2007). Therefore, in the conventional system, the space–time continuum information of tissue injury is destroyed because it decomposes into a cell and measured. The structure is also destroyed if leukocytes from multi-tissue injury are mixed by the observer. Both cause the destruction of the structure, especially statistical sampling. In modern pathology, the CET faces problems. However, the leukocyte continuum (LC) as SτC as RDE, which realises the SOC, solves the problems of structural bioinformatics and enables the measurement of local inflammation and tissue information. The problems encountered are 1) lack of structure of bio-information, which can measure the space of inflammation over time {problem of cell/tissue injury (cell/tissue information) }; 2) destruction of the structure of bio-information by statistical sampling; 3) lack of smoothness of differentiation; 4) problem of the simultaneous differential equation (SDE) (prey and predator, etc.); and 5) the problem that cannot be visually quantified by a doctor.

## SYSTEM AND METHODS

### Materials

In my clinic, the LC was based on the finding that was incidentally discovered in the microscopic examination of periodontitis (infection). Microbial culture and microscopic examination to the infection were also performed in the clinic. {I have posted using the finding for journal, and such as society, on the noticeboard of my clinic for progress of medical science (Text S5).}

The LC had the SOC as the material. The SOC was built as a continuum formation by water treatment of leukocyte(s) as an arithmetic circuit material. The RDE was the method used to read the LC. Meanwhile, kOOC (OOC with Kyoku), zOOC (OOC with Zai) and kzOOC (OOC with Kyoku and Zai) were the solutions used for the SOC. The RDE (Definition S6) is composed of Zai and Kyoku, which was a primitive-operator (Definitions S1–6). In the microscopic examination, a multicircle explorer (MCE; Isizuka Co., Ltd., Japan; Figure S1) was used to sample the gingival crevicular fluid (GCF) using glass slides, cover glasses, a phase-contrast microscope and a fluorescence microscope (Corporation Riken, Japan). The OOC used a general-purpose PC, picture processing software and arithmetic software.

### Methods

#### Conservative sampling, water operation and generation of the WSTLs/WTLs

The GCF from the periodontal pocket of patients with periodontitis was conservatively sampled using an MCE (Figure S1) and transferred to glass slides. Accordingly, 20 µL of water (water treatment of leukocyte: WTL) and 20 μL of acid red aqueous solution (0.1 wt.%) were added (water solution treatment leukocyte: WSTL). The samples were covered with glass cover slips and observed using microscope images {D()} (size: 720 × 480).

Through the water treatment, the mutual leukocytes combined together for a short distance (Figures 1 and S4–S6) and formed the LC, which had SτC and τSC. Therefore, the water treatment can prevent the destruction of the space–time structure of the leukocytes from the tissue injury and the destruction by mixing of the leukocytes from the multi tissue-injury. The size was assumed to be proportional to that of the tissue injury because larger leukocytes exist in larger injuries. The larger cluster that exceeds that of a small tissue injury does not exist (Figures 1–5 and S2–7).

**Figure 1.**
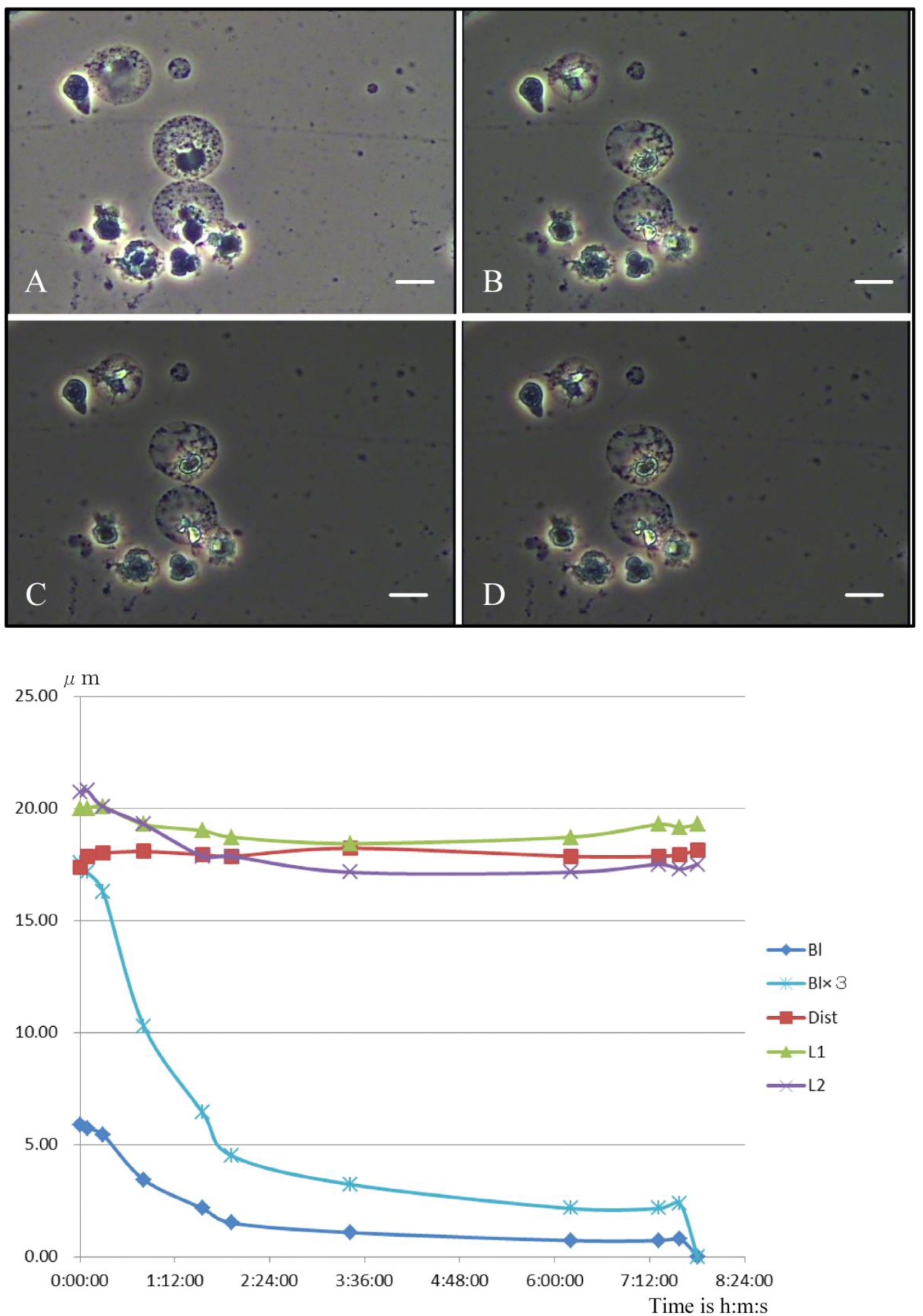
Leukocyte–dissociation curves. **(A)** Time 0:00:00 is the start of the experiment. The contact line (Figure S2) represents a primary bond. **(B)** Time 6:12:10 is the point after initial dissociation. Coupling occurs via a secondary bond between two leukocytes. **(C)** Time 7:48:23 is the point of secondary dissociation and bond breakage. **(D)** Time 8:01:37 is the point after secondary dissociation. (**A**–**D)** Scale: 10 μm. **(E)** The bond length (Bl) describes the length of the contact line formed by the primary and secondary bonds. Leukocyte 1 (L1) represents the upper leukocyte diameter, whilst L2 represents the lower leukocyte diameter. The distance between L1 and L2 is denoted by ‘Dist’.

#### Time and phase condition of the WSTLs/WTLs

The active leukocytes (ALs; Figure 2), destructive leukocytes (DLs; Figure 3) and LC are presented in Definition S7 (Figures 2–5).

**Figure 2.**
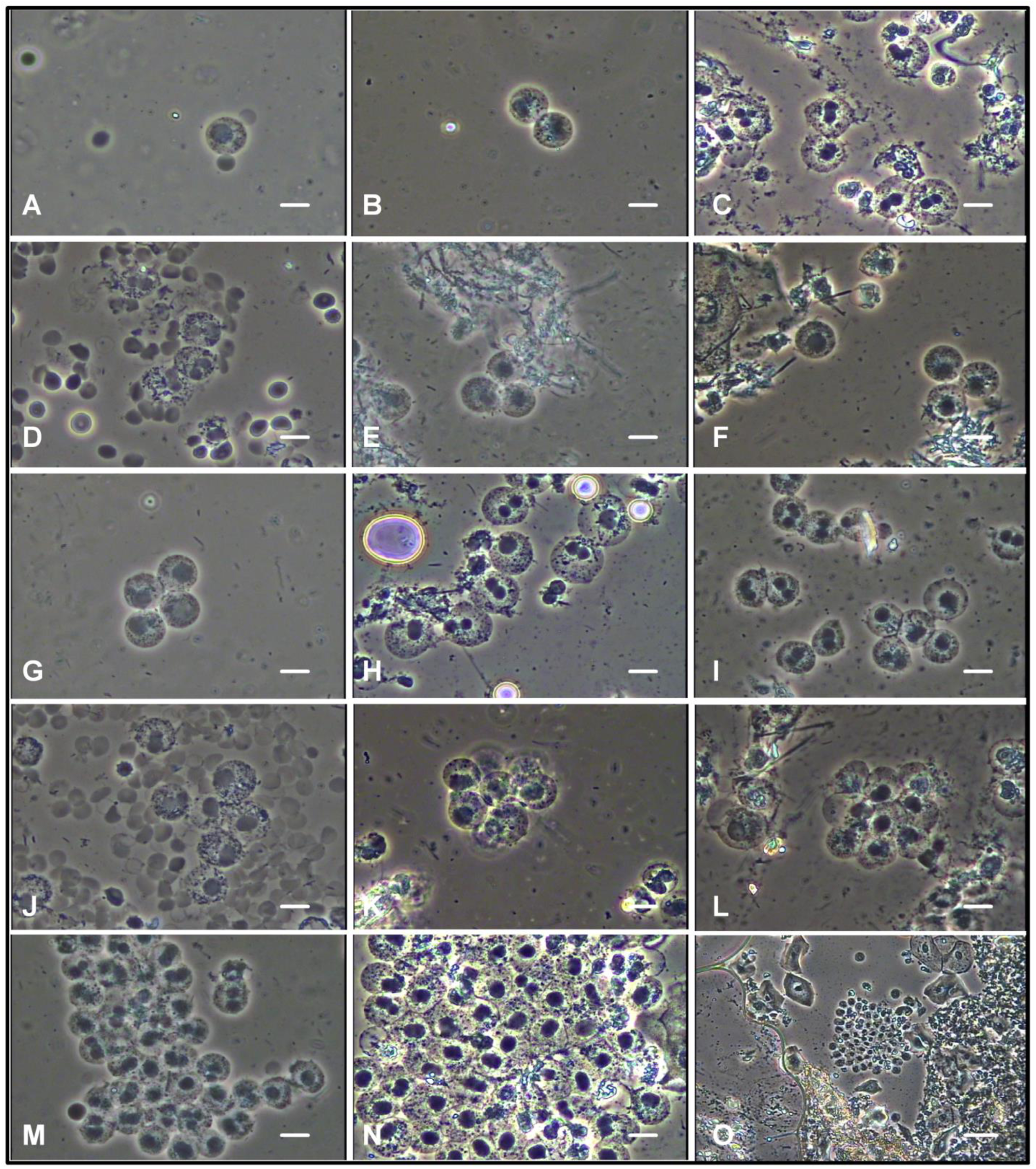
AL clusters identified according to the dynamic movement of the organelles following WTL. **(A)** Single and **(B–O)** multiple leukocyte clusters **(A–N)** Scale: 10 μm and **(O)** 40 μm. Before applying the RDE, ‘Model’ is seen as the CDE with many singularities. It is perceived as SOC after RDE application.

**Figure 3.**
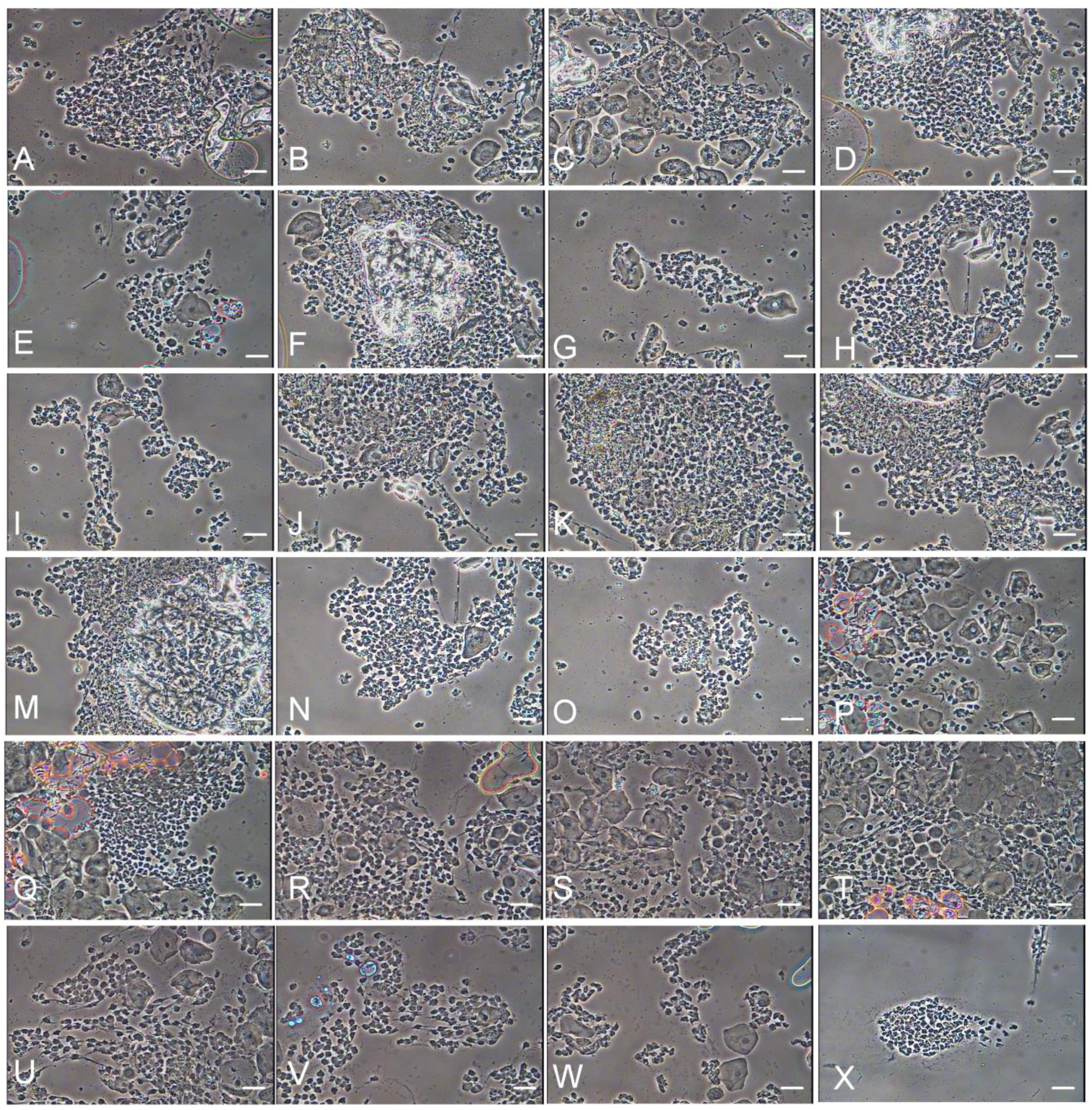
DL clusters. Destructive leukocytes (DL) clusters. Scale: 40 μm. DL cannot be analysed by CDE. The analysis is possible for SOC. However, DL gradually loses continuity. SOC also collapses according to it. However, DL is useful to diagnostics.

#### SOC and OOC observation

Reading the SOC image is possible with the RDE (science eye) in the brain (i.e. the SOC recognises the graph as a result of arithmetic, operation and calculation) (Definitions S1–7, Method S1). The OOC was calculated using a common digital computer.

### Algorithm

#### First algorithm of the arithmetic circuit creation

##### Circuit elements

A conventional computer consists of AND, OR and NOT gates in a logical circuit. This leukocyte circuit (LC) as SτC consists of Zai and Kyoku (Definitions S1–S3). The WSTLs/WTLs are Zai, and had an inner-space and inner-time. The inner-space herein is Δ*N* (zero dimensional space; Po is the number described in Definitions S1, S3, and S7). The inner-time is Δτ (Po is the fixed-time described in Definitions S1, S3 and S7). ZaiN and Zaiτ (Δ*N*/Δ*τ*) consist of WSTLs/WTLs (Figures S4–7). ‘*N*’ is the number of cells {leukocyte(s) or microorganism(s)}. t_n_ is the physical time. *τ* denotes the phase time by a biological activity time (Figure S8, Definitions S1.6 and S7). The changes in the boundaries (as the membranes of cells or nuclei) are the changes of the phase time (the main value of *τ* is ‘1’; *τ* = 1). The boundary defines the inner-Kyoku (iK; Definitions S1–S3). Zai constitutes a real coordinate system (RCS; Figure 6 C3-RCS). C1 RCS and C2 RCS are the components, which did an inheritance derivation from C3 RCS (Definitions S3.6.1 and S6.1 and Figure 6). n and m are the phase values as positive integers (Definition S1). n is origin (relative or absolute), whilst m is non-origin.

**Figure 6.**
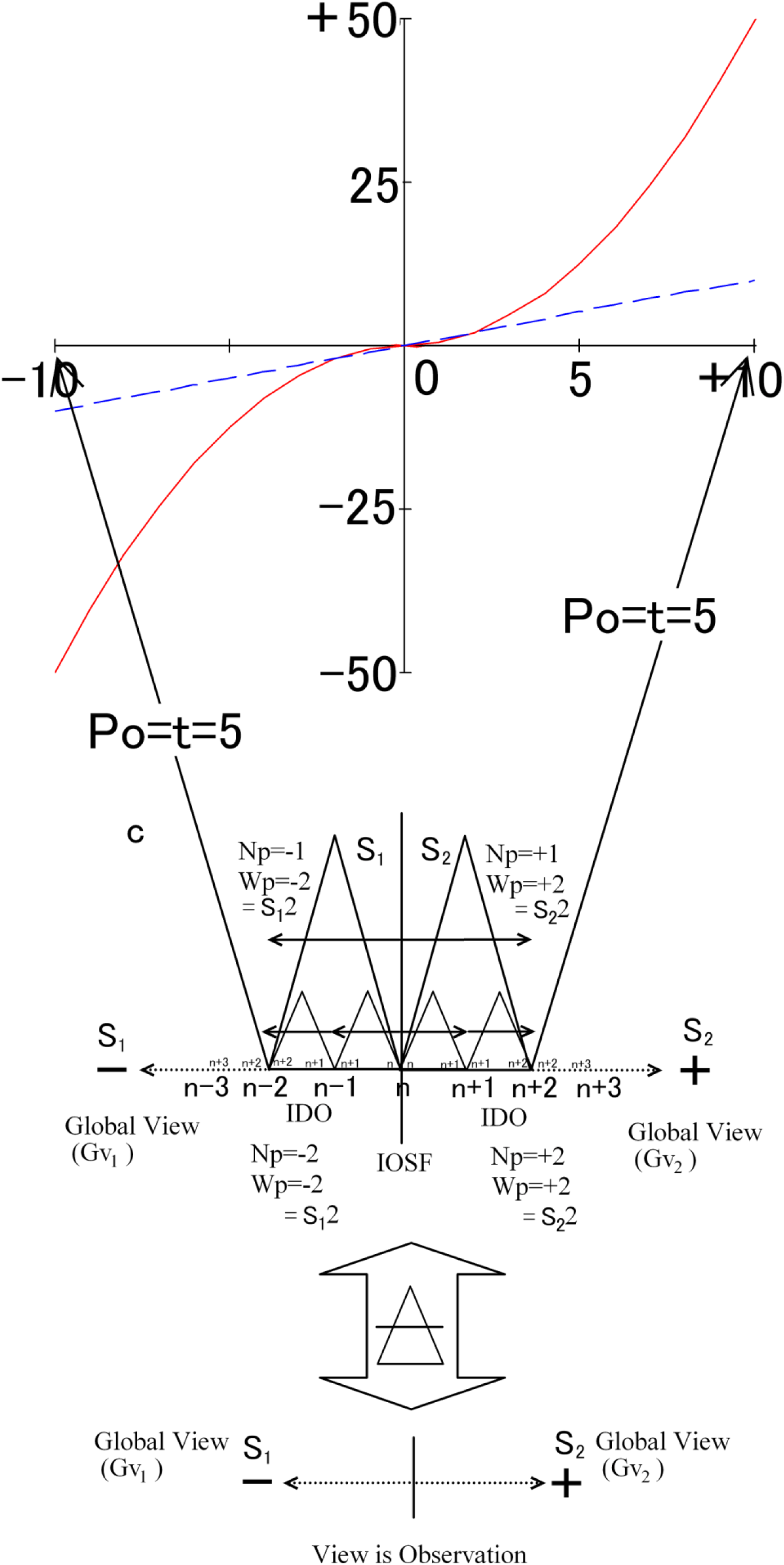
Real coordinate system. Condition 3, coordinate system according to StrZai on the G-axis (horizontal axis). The orthogonal axis is the K-axis. Single StrZai (Np = S_1_·1 = −1, Wp = S_1_·2 = −2) (Np = S_2_·1 = +1, Wp = S_2_·2 = +2) and StrZai continuum (Np = S_1_·2 = −2, Wp = S_1_·2 = −2) (Np = S_2_·2 = +2, Wp = S_2_·2 = +2). Po is time (*t* = 5).

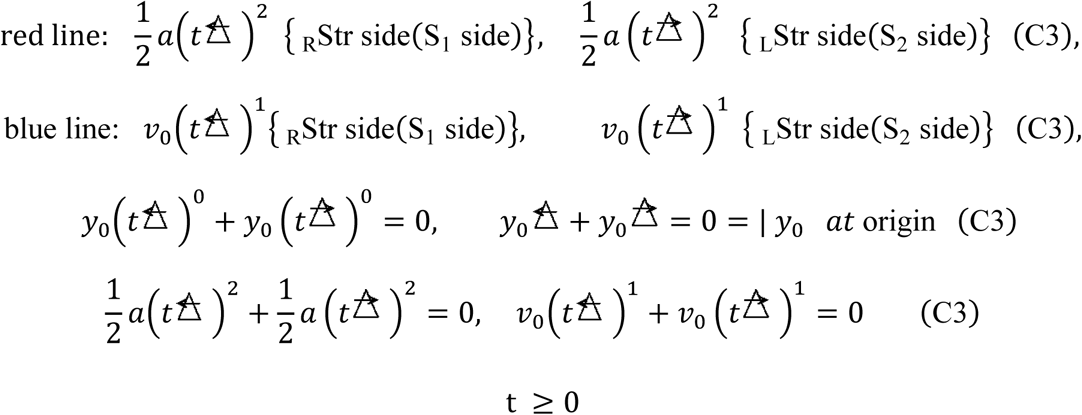

The C3 coordinate system solves the problems of singularity, integration constant and initial value. The red (and blue) line denotes the ‘right line’.

##### Circuit build

Considering whether leukocytes were immersed in water and near tissue injury, the leukocytes were conservatively sampled by the MCE (Figure S1) and immediately immersed in water or water solution (Figures 1, S2, and S3). The microscope images of these leukocytes were then used to generate single-layer leukocytes (SLLs; Figures 2 and 3), string LC (StrLC; Figure 4) and multilayer LC (MLC; Figure 5). These were equivalent to the arithmetic circuit itself, and could be used as the RDE (Definitions S1–S7 and Figures S4–S7). In other words, the equation and the operator(s) that elementised the nature formed the structure of the arithmetic circuit and the resulting graph. The structure forms the arithmetic circuit and the resulting graph if the natural phenomenon is constructed and built on certain conditions. Accordingly, it defined the SOC.

**Figure 4.**
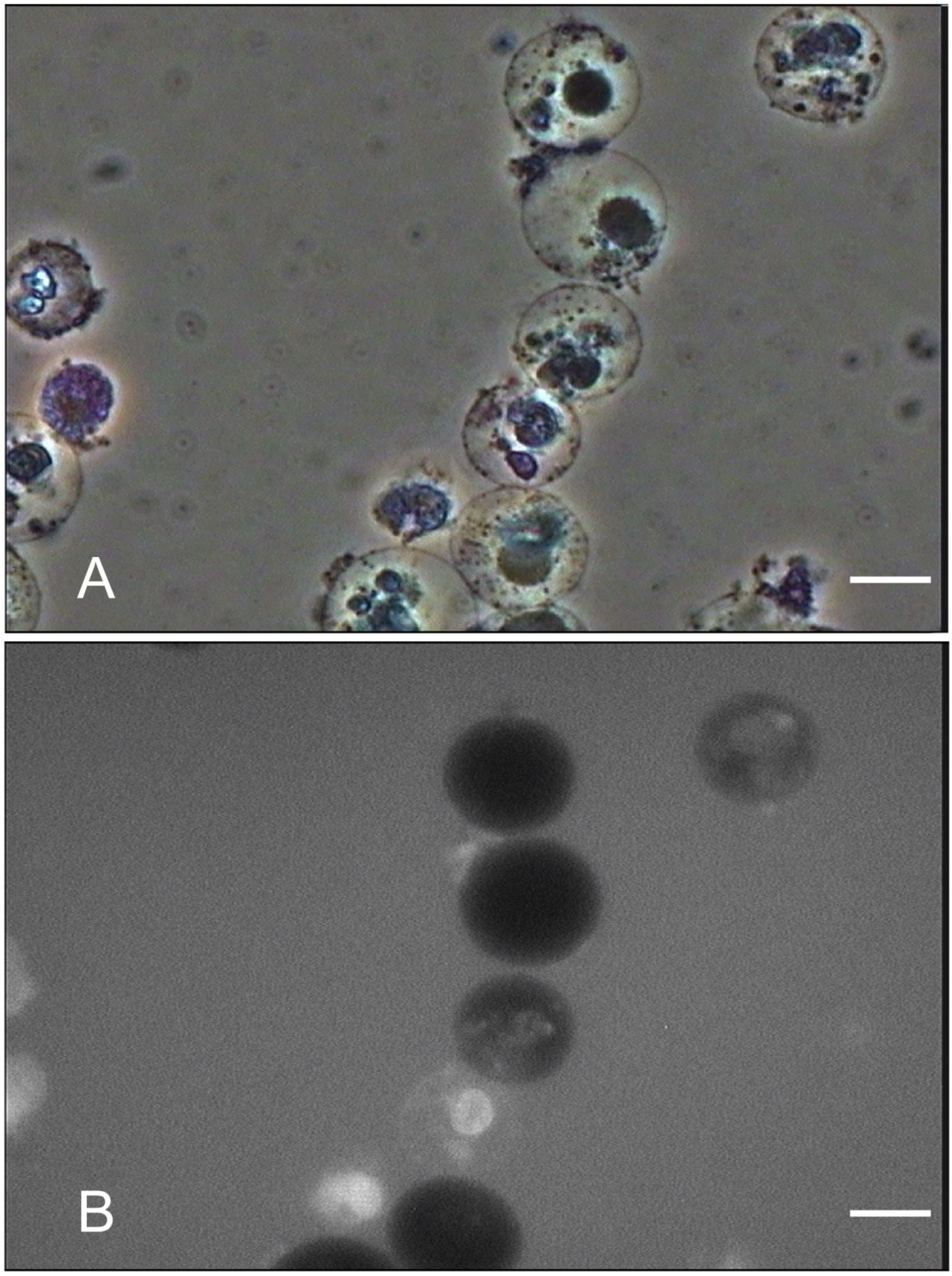
StrLC by WSTL. Results of the WSTL as measured by fluorescence. **(A)** Results of the WSTL according to a phase-contrast observation. Older cells are white, whilst newer leukocytes are black. Scale: 10 μm. **(B)** Results of the WSTL as measured by a fluorescence mode observation. Older leukocytes are white, whilst newer leukocytes are black. Scale: 10 μm. StrLC is a string function with a CS as an independent axis. AO-or RO-type singularity is observed according to this function. StrLC shows us various problems, such as EOM and CET. The calculation of AO or RO is possible using primitive-operator.

**Figure 5.**
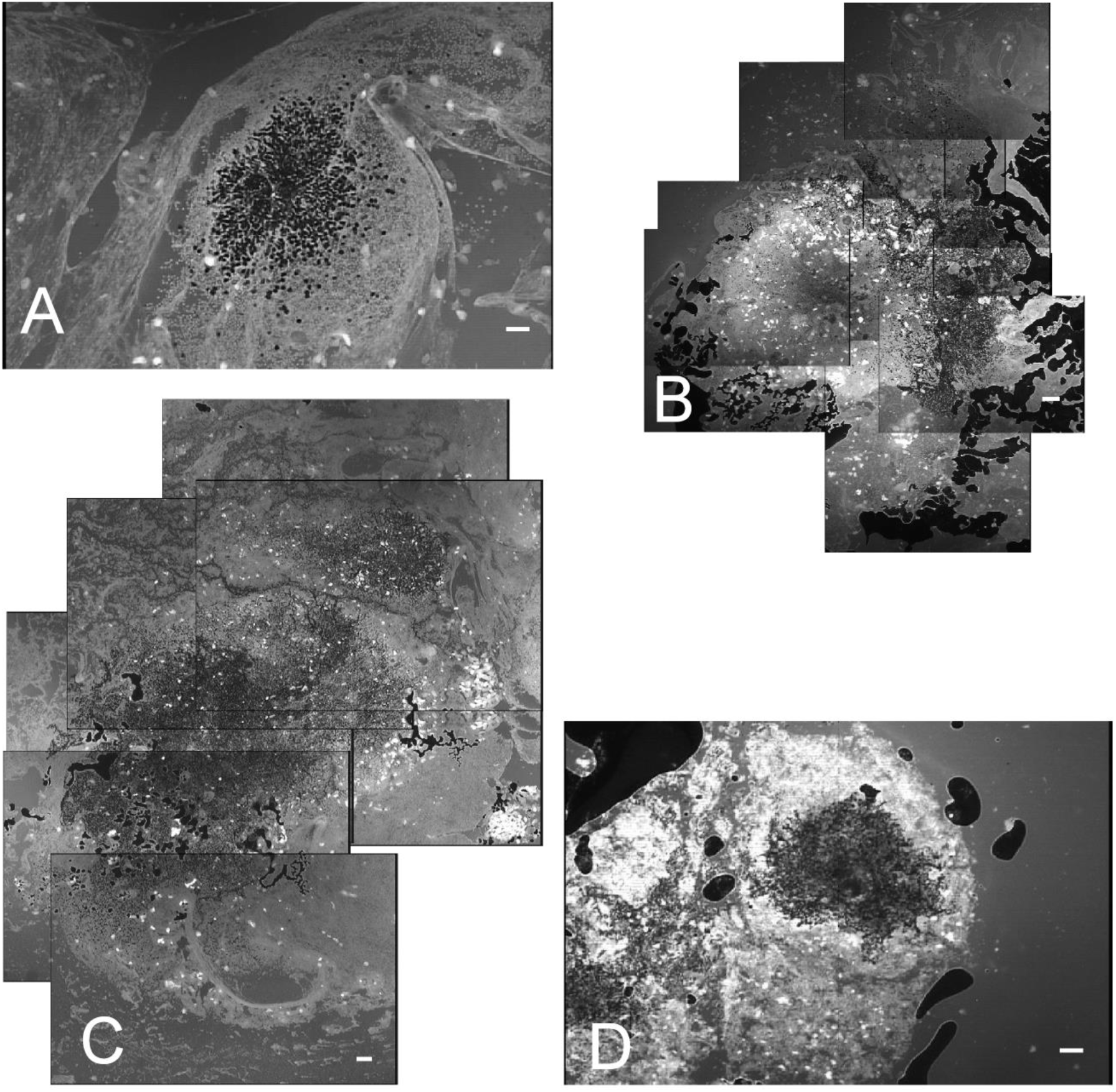
MLCs as SτC by WSTL. An AO-type singularity is observed at the centre of the MLC. **(A)** Enclosure at the outermost periphery of a fibrous connective tissue indicating a complete view of one tissue injury. Scale **(A, C and D)**: 100 μm and **(B)** 200 μm. Before applying RDE, ‘Model’ is perceived as the CDE with many singularities. After applying the RDE, it is perceived as SOC. It has solution as OOC. MLC shows the complete SOC RDE. MLC can observe gsOOC simultaneously.

The LC in the SOC is defined as SτC. The LC in the OOC is defined as τSC (Definition S7.2, Figure S7).

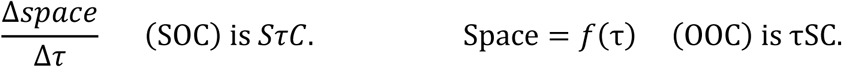

The SOC denotes the microscope images that simultaneously provide information on size and form and represent an RDE. In contrast, the OOC is the solution.

### Various LCs as the SOC as the arithmetic circuit and OOC

#### StrLC

##### SOC

The StLC is the string LC, whose width ∅ is one WSTL/WTL. Therefore, the size of the tissue injury, which it indicates, is ∅ (Figure 4, Table S3). Therefore, t_n_ denotes the physical time.

##### SOC1 (from C1-RDE)

The injury size is (*N* = 1), where *N* is fixed. Therefore, the equation uses time as a variable:

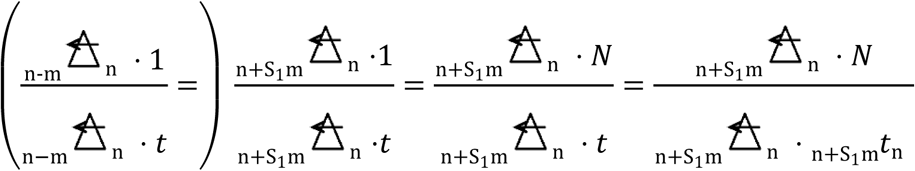

A motion object without iK: DL (AL → DL):

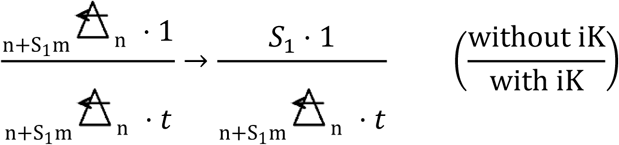

S_1_ is defined in Definition S2.

##### SOC2

Tissue injury form: 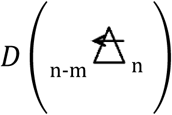, equal to one WSTL/WTL circle form You will observe StrLC as 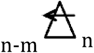 in the microscopic image in 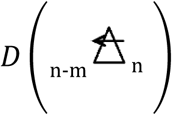.

##### OOC

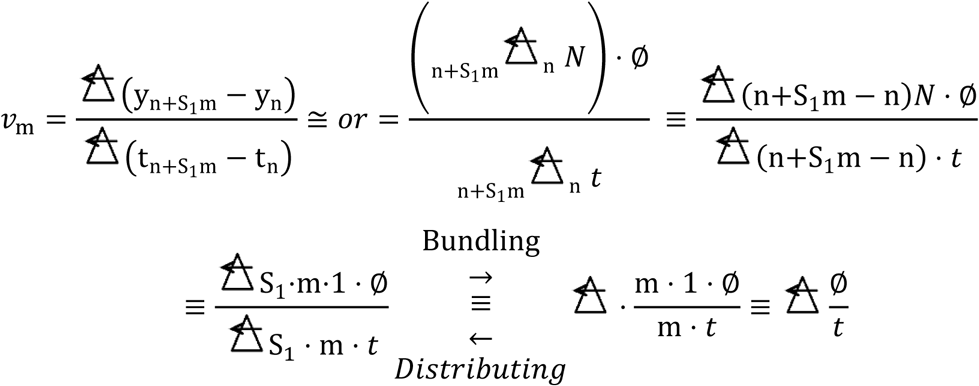

In StrLC, *N* = 1, n is origin (AO or RO), and ∅ is one leukocyte diameter.

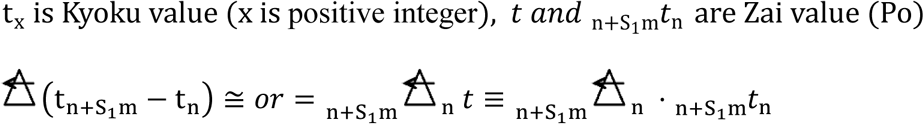

### Properties

A StrLC from the same injury can be described by the EOM; however, this StrLC biomarker uses the RCS to represent future inflammation, making it necessary to solve the problem of IDP-EOM by addressing smoothness. A cOOC expresses only one side. Therefore, it generates singularity and integration constants (Figure S9, Table S1). The EOM with Str Zai C3-CS (CS: coordinate system) (Definition S3.6.1, Figure 6, Text S6) is presented as follows:

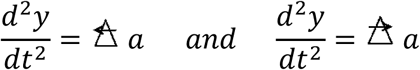

This is only a ‘model’.

The OOC with iK are

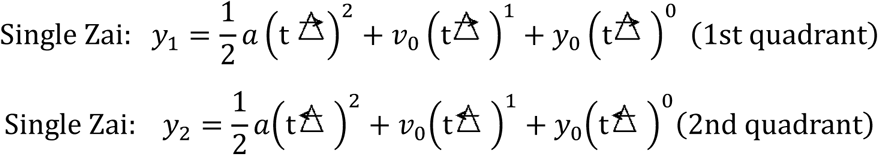

Above the single Zai is the ωd switch that is ‘ON’. In this case only, t is a variable. The other is Po (Supplementary Method S1.1). The others are all ‘OFF’. The variable is phase (n and m). Zai continuum:

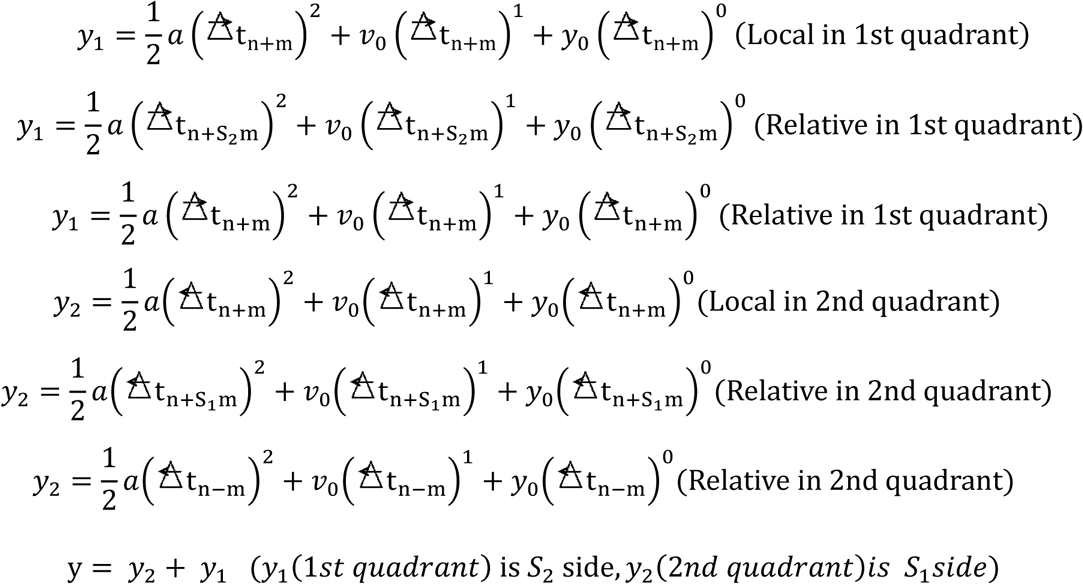

*a* is the acceleration as Po; *v*_0_ is the velocity as Po; and *y*_0_ is the distance as Po. *S*_1_ and *S*_2_ are the Str operators (Definitions S2 and S3), and the observer in the local field (LF) is denoted as n + m. The operation was performed using the LF and the relative field (RF). The observer in the global field (GF) RF is denoted as n + S_1_m in the C3-CS, where the independent variables included 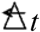. n is the absolute or relative origin. Conventionally, at an origin, the direction of motion of the quadratic term is reversed (Figure S9). However, in an RCS, the quadratic term maintains the same direction, and the linear term of the conventional equation and of this equation maintain the same direction (Figures 6 and S10, Table S1).

*y*_*x*_ is the diameter ∅ of the WSTL/WTLs, whilst *τ*_*x*_ is calculated from the activity of the leukocytes. When *y*_1_ = ∅_1_, the velocity is calculated from the solution of the EOM {RF (real field)}:

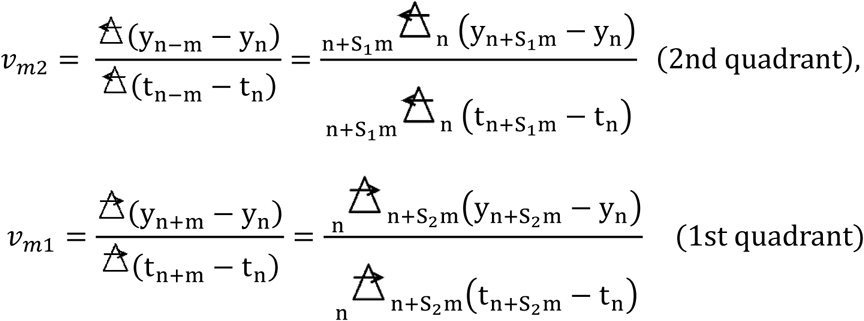

The object without iK (time has iK) is presented as follows:

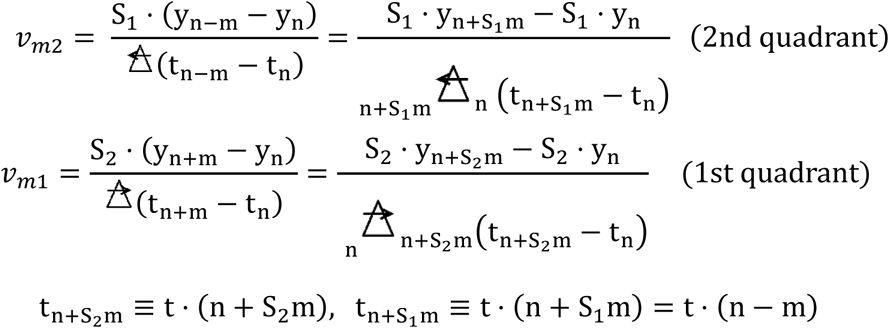

y_1_, y_2_,…y_n_ is one WSTL diameter, and t_1_, t_2_,…t_n_ is the elapsed time of the WSTL. Acceleration is calculated from the solution of the EOM {RF (real field)}:

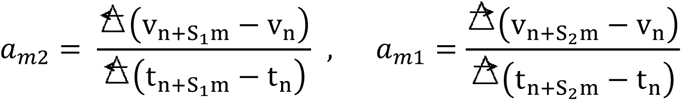

where v_1_, v_2_,…v_n_ denote velocity, and t_1_, t_2_,…t_n_ denote the elapsed time of the WSTLs.

### SLLs (AL or DL)

C1-RDE (Definition S6):

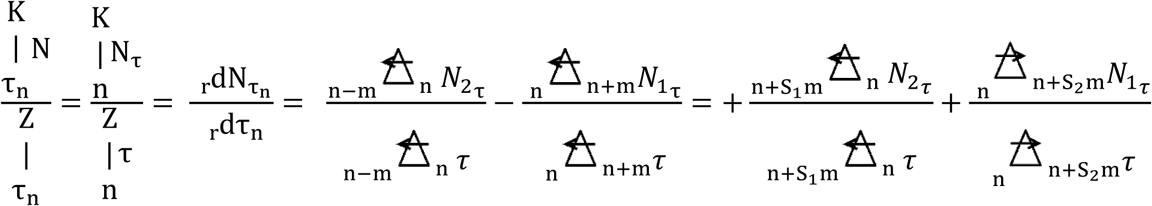

Detail:

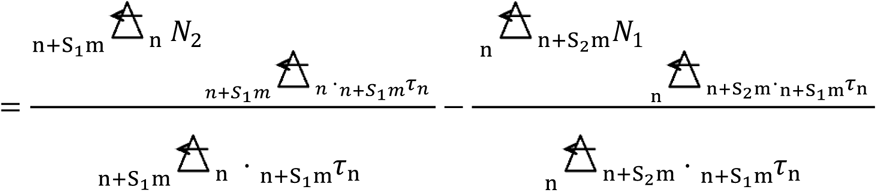

### Properties

The cases in which one *N* in C1-RDE is zero (or unknown) are called SLLs. The unknown cases cannot be calculated. However, if one term is zero, the SLLs are equivalent to C1-RDE (MLC); ALs are the acute stage (*N*_2_ > 0 and *N*_1_ = 0); and DLs denote the improved stage (*N*_2_ = 0 and 0 < *N*_1_). In the unknown cases, the ALs indicate the present inflammation, whilst the DLs indicate the past inflammation. Even if the other terms present the kind of values, the size of the LC is larger if the injury is larger.

### SOC of the SLLs

The SOC of the ALs (Figure 2) is the right first term of the RDE, and the other term is zero. The SOC of the DLs (Figure 3) is the right second term of the RDE. The tissue injury size is nearly equal to the size of the WSTL/WTL cluster (Section **MLC**). The SOC evaluates the SLLs (leukocyte cluster with the same *τ*, Figures 2 and 3), where *N* is the number, and *τ* is the fixed time.

An example of the ALs is (Figure 2, from AL0 in Figures 5A and S5–7):

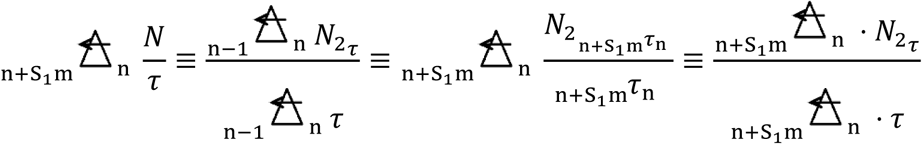

An example of the DLs from AL1 is (Figure 3, from AL1 in Figures 5A and S5–7):

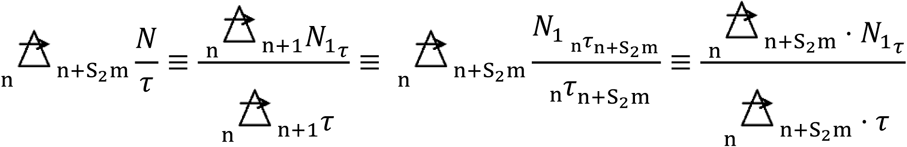

AL → DL (life → non-life):

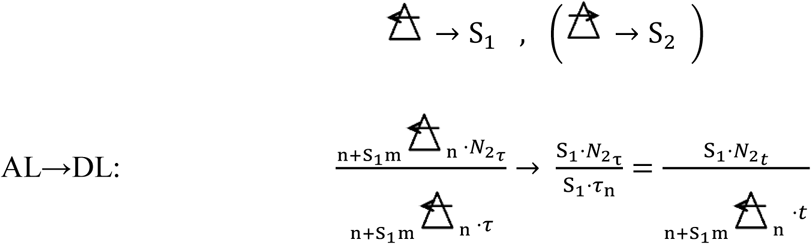

In the case of the DL (Figure 3),

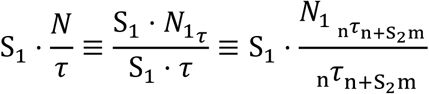

where a single value of *m* = 1 exists; *τ* is fixed time; *t* is real time; AL represents the active leukocytes exhibiting a dynamic movement (Figure 2); and DL does not have a dynamic movement (Figure 3). *N* is the number of leukocytes. *τ*_n + m_ is the phase time.

In the case of the AL (*τ* = 1) (Figure 2):

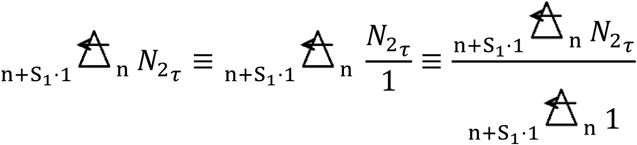

In the case of the DL (Figure 3):

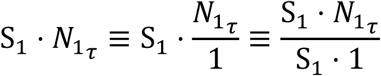

The SLL has the same τ value (*m* = 1), and the size of the SτC represents the size of the injury.

### MLC

#### SOC

The SOC shows the size of the injury as an MLC:

The newer side term is 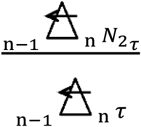, whilst the older side term is 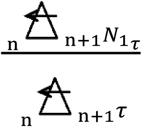

In detail,

Newer side term: 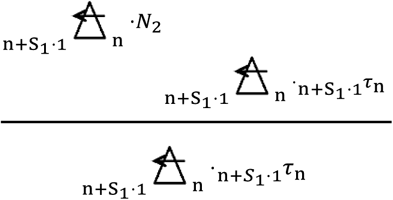

Older side term: 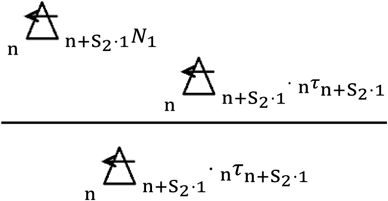

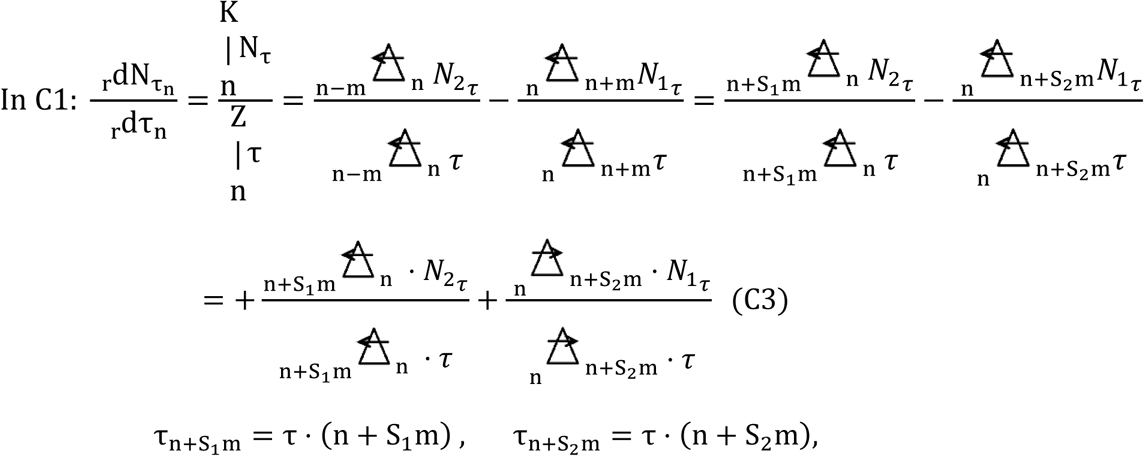

The numbers of leukocytes *N*_2_ and *N*_1_ and 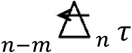 and 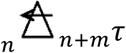 as the phase time are extracted from the LC as SτC in Figure 5A by a biological activity time chart by leukocytes (Figure S8) (example of the time is “τ = 1, 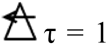 in LF, and 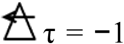 in RF”; Definition S3). When a newer side term (*m* = 1) represents a core leukocyte, the size of the injury is represented by this as the LC (Table S2). SτC is the same as a CS that is equivalent to deform the dependent-variable-axis of the orthogonal-CS (Figure 6).

### Fluctuation velocity (Kyoku) (***γ*N)**

The fluctuation in the velocity is described by C1-RDE:

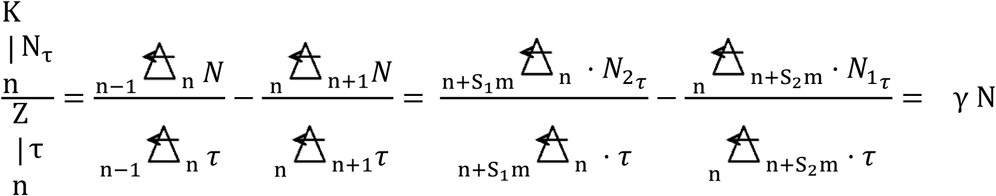

The inflammatory condition of each area of the *τ* zone is given as follows:

1. An older zone being the same size as a newer zone represents a chronic stage: {*N*_2_ = *N*_1_ (each *N* ≠ 0)}.
2. If a newer zone is smaller than an older zone, inflammation is in the improving stage: (*N*_2_ < *N*_1_).
3. If a newer zone is larger than an older zone, the inflammation progresses to an acute stage (*N*_2_> *N*_1_).
4. *γ*N is visualized (SOC) (Definition S6.2).

*γ in OOC (in C1, C2 and C3 RDEs)*

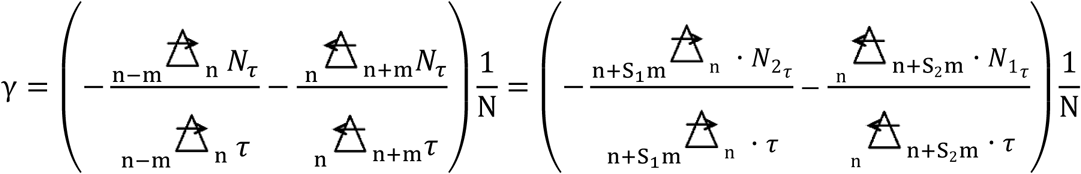

The usual absolute value operator is as follows:

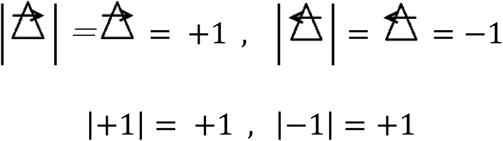

The relative absolute value operator (relative absolute operator):

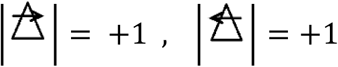

The relative absolute value is as follows:

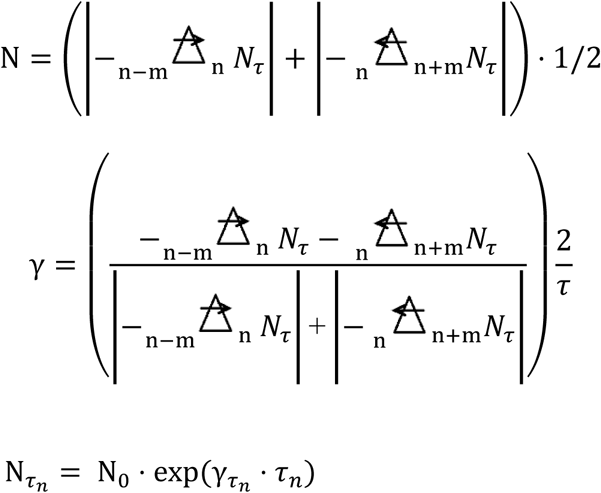

N_0_ is the initial value; γ = 0 is the chronic stage (each *N* ≠ 0); γ < 0 in C1 is the acute stage; γ > 0 in C1 is the improving stage (RF); γ < 0 in C2 is the improving stage; and γ > 0 in C2 is the acute stage (RF and LF).

### γ of SLLs

The SLLs cannot be used to obtain the solution if there is no other term. In contrast, γ = 0 is the chronic stage if there is a term (the term is zero). C1-RDE: γ = S_1_(+2) is the acute stage (*N*_2_ > 0, *N*_1_ = 0); γ = S_1_(−2) is the improving stage (*N*_2_ = 0, 0 < *N*_1_; RF); C2-RDE: γ = S_2_(−2) is the improving stage (*N*_2_ > 0, *N*_1_ = 0); and γ = S_2_ (+2) is the acute stage (*N*_2_ = 0, 0 < *N*_1_; RF and LF).

### Antigen antibody reaction (background is Text S7)

The fluctuation of γ according to an antigen–antibody reaction is presented as follows (C4, Definition S6):

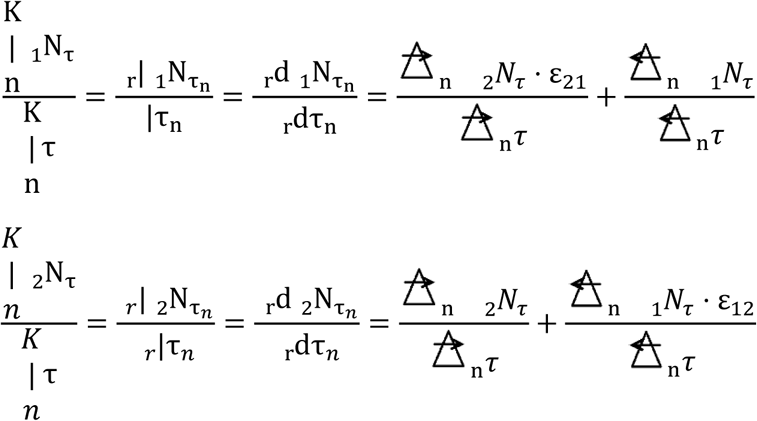

_1_*N* depicts the leukocytes; _2_*N* indicates the infective antigen (bacteria or microorganisms); ε is an interference coefficient; ε_12_ is the phagocyte number coefficient pertaining to the leukocytes; and ε_21_ is an invocation (cytokine) coefficient of the leukocytes to an antigen.

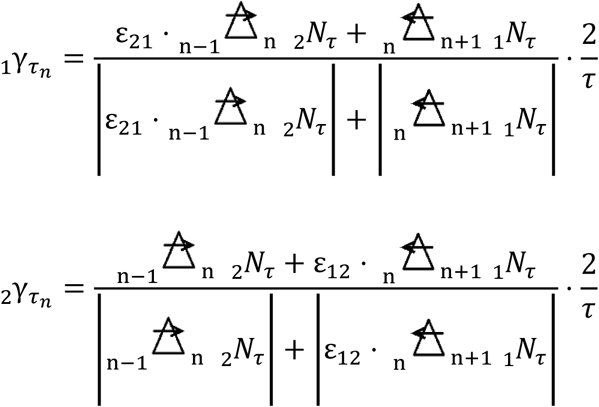

_1_N_0_ is the initial value of leukocyte, and _2_N_0_ is the initial value of bacteria:

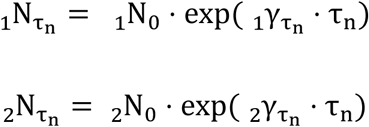

The *τ* ratio (τ_r_) is used if the reaction time of existence 1 (_1_N) and existence 2 (_2_N) is different:

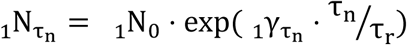

As a result, a stable solution is obtained (Figure 7). The conventional SDE was unstable by above different-object interference (AO), and the solution described vibration and rotation, where the new SDE successfully converges. Therefore, the future of inflammation becomes clear.

**Figure 7.**
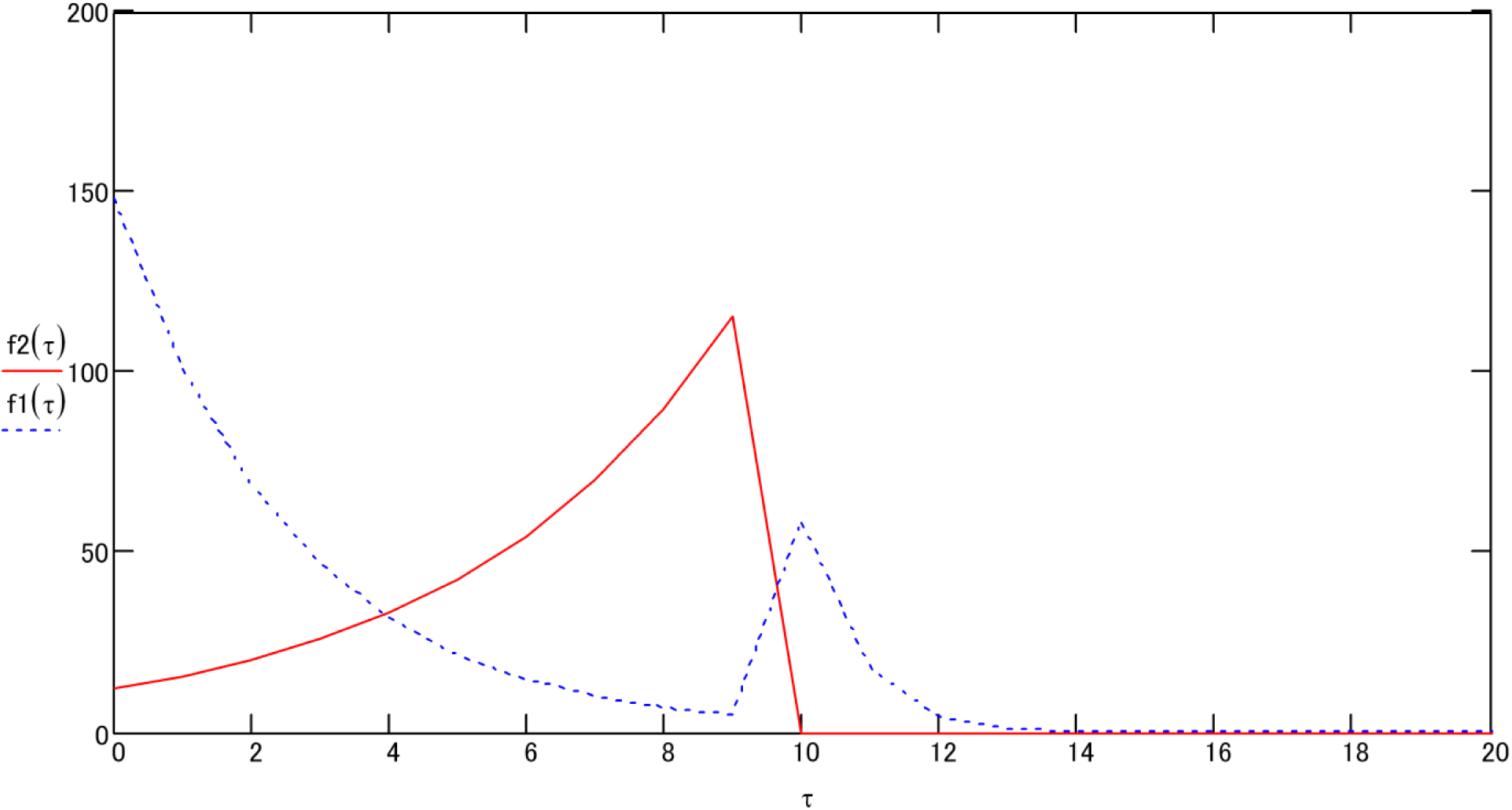
Antigen–antibody reaction according to Lotka–Volterra equations. The solid line represents bacteria, whilst the dashed line represents leukocytes.

## IMPLEMENTATION

### StrLC

The SOC simultaneously presents size and form. Time is calculated by the OOC; hence, the SOC is inaccurate.

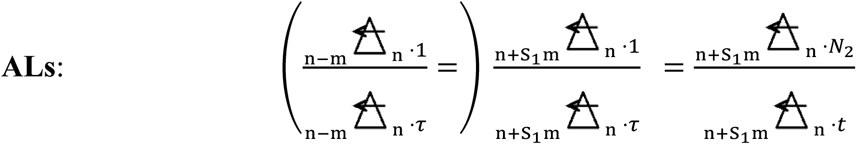

*t*: physical time

Size:

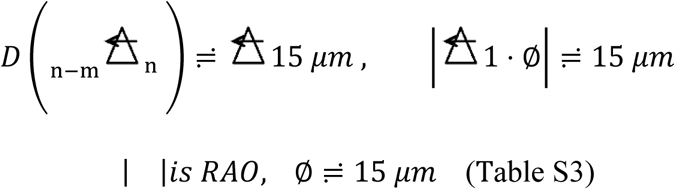

The tissue injury size is 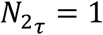. The size is the same as WSTL/WTL (Figures 2, 4 and S4 B, C, D, and H). The space information was constant, whilst the time information was variable.

Therefore, the form of the injury predicted from this StrLC form is the injury of a pipe-like form of approximately 15 μm in diameter.

### OOC

The transition from stage AL1s to AL2s (Figure S8) took 28 min (Table S4). (m = 1, *N* = 1)

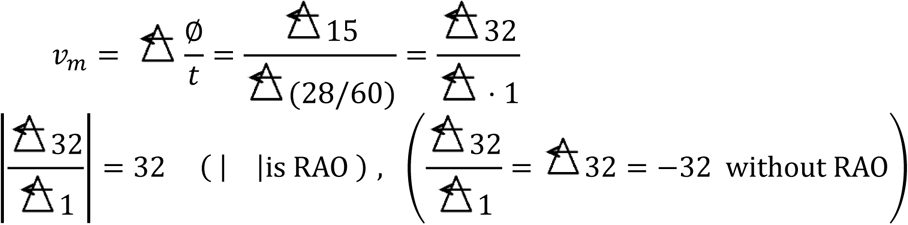

Accordingly, ∅ = 15 μm (Table S3). The velocity is 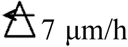. The tissue injury size is 15 μm (constant). The transition from stage AL0s to AL1s took 4 h 24 min (264 min, Table S4) at 3.4 μm/h.

### Conventional problem observed from StrLC SOC and OOC

The dynamic information for this StrLC included time and cell injury information. The tissue injury information was constant. Using StrLC, we could understand the problem of observation via a one-cell unit, which enabled us to elucidate the problem of Lcnt and FCM.

### SLL

#### SOC

*τ* is the phase time in Figure 2 (τ = 1). The SOC can be located with SOC1, SOC2 and SOC3 in order of thinking. You will realise the size of inflammation and the future stage by the SOCs. **SOC1**: Injury form

A figure (graph) of 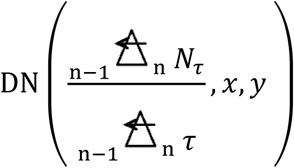 is in each envelope (white closed line) in Figure S4. *x* and *y* denote the microscope images on a 720 × 480-pixel orthogonal-CS. The other variables are the same as the SOC3.

### SOC2

Injury size

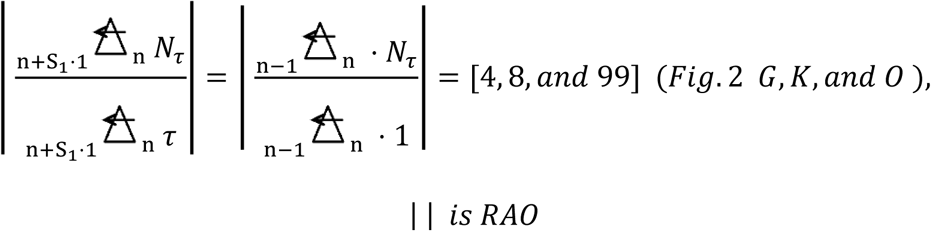

### SOC3

Graph of size and form

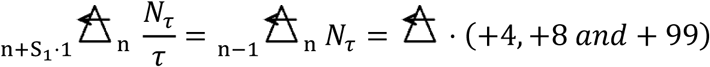

is the acute stage. The injury size is the SOC 2. The injury form is SOC1.

### OOC as a part of C1-RDE (MLC)

If the other term is zero, γ = −2 = S_1_2 is the acute stage (*N*_2_ > 0, *N*_1_ = 0; Figure 2). Figure 3 shows the improving stage (γ = +2 = S_2_2; *N*_2_ = 0, *N*_1_ > 0).

### MLC

**SOC** (Applied data: Figures 5A and S5–S7, Table S2)

**SOC 1** Injury form:

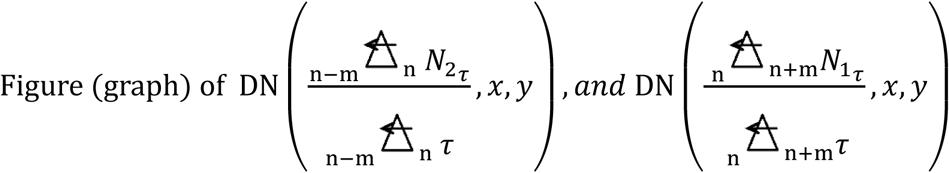

based on the insets (white closed line) in Figure S6.

The upper term is the number of leukocytes (size), whilst the lower term denotes a certain *τ* zone of one leukocyte cluster.

**SOC 2**: Injury size (Table S2)

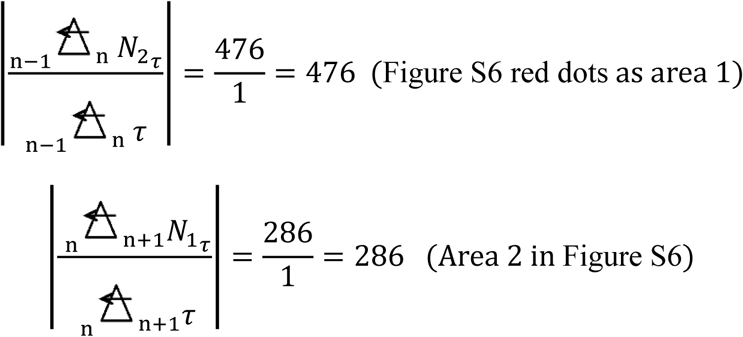

τ =1, | | is RAO; the core is area 1 (red dots); and outside is area 2.

**SOC 3**: graph of size and form injury:

### Area 1 (AL core)

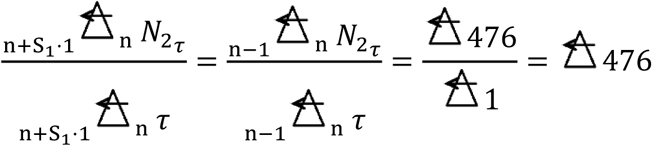

#### Area 2

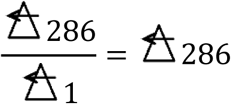

The figure (graph) is the result of the RDE of areas 1 and 2, and one may be able to count in a moment in this graph (as **SOC1-3**).

**SOC 4**: Fluctuation velocity and *γ*N; Kyoku

From Table S2 (*N*_2_ = 476, *N*_1_ = 286):

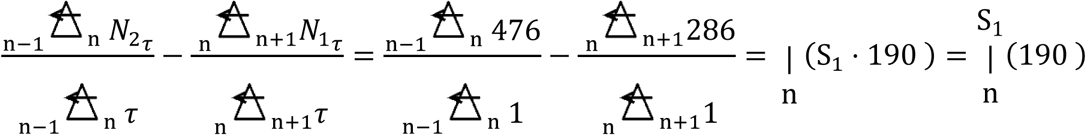

The fluctuation velocity value (*γ*N) of Kyoku is {|(S_1_190)}, representing the acute stage. *γ*N is visualized (SOC).

### OOC

Table S2 shows that γ was |(−0.5) = |(S_1_0.5) in C1 (RF), representing the acute stage.

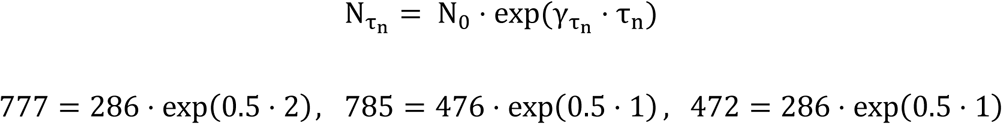

S_1_ is the abbreviation. This equation is discovered as gsOOC in a part of SOC LC in Figures 5A and S7, and will be presented as a bar graph or a line graph. Naturally, γ of an antigen is required.

### Antigen–antibody reaction (prey and new predator) (C4-CS)

#### SOC (Figures S11 and S12)

The display of the number of bacteria in a certain *τ* time and the number of leukocytes in the *τ* time were pseudo-SOC because ‘ε’ could not be used for the SOC. Moreover, the number of bacteria was not accurate in this case. Therefore, the SOC was also inaccurate. In the future, the display of SOC must be improved, and ‘ε’ must be used for the SOC.

#### OOC (Figure 7)

The solid lines depict the bacteria, whilst the dashed lines depict the leukocytes.

As an example, γ_b_ = |S_1_0.50 (Table S2, S1_Supporting Information); γ_g_ = |S_2_0.25 (Marsh et al., 1994); ε_21_ = 0.5 (ε_21_: arbitrary numbers were substituted; the appropriate number will be determined in future studies); ε_12_ = 250 (Phagocytosis Assay Kit, https://www.funakoshi.co.jp/contens/3695); f_2_(τ) is the number of bacteria; and f_1_(τ) is the number of leukocytes. The relative τ ratio (τ_r_) is 1.3 (1:1.3; τ_r_: arbitrary numbers were substituted; the appropriate number will be determined in future studies). Accordingly, τ = 10 at the time of interfere. The number of antigens was set to 0 when the antigen was set to 0.

The dissolution of the AO and the RO singularity using view-operator was successful for the OOC.

## DISCUSSION

As mentioned earlier, the RDE is put in the brain. When observing Figures 2 to 5 and Figures S4 to S7, one will recognise having become Dr. Perio (specialist periodontist), with the Perio eye (periodontal-diagnostic doctor’s eye) by the SOC and gsOOC. Accordingly, the SOC has a real-time property. No ‘model’ exists, and no complicated property, like OOC, calculates the solution. The SOC will educate A.I. and the young doctor. Furthermore, the OOC from the SOC will extend it more precisely.

Conventional science only has the OOC; therefore, science has various problems. The problems of cOOC are also mostly shown by the SOC, and will mostly be solved by the SOC. At such time, the LC was found as the SOC in the periodontitis examination at the clinic. An example is the inflammation, in which cell information from cOOC has problems. Therefore, inflammation is described by the SOC and a new OOC from the SOC. Moreover, the LC as the RDE, which realises the SOC, solves the problems of structural bioinformatics and enables the measurement of local inflammation:

1. The SOC (LC) has the structure of bio-information, which can measure the space of the inflammation in time. Therefore, it solves the problem of cell injury (cell information) and tissue injury (tissue information).
2. The SOC LC exhibits no destruction of the structure of bio-information by conservative sampling.
3. The SOC as LC by the RDE has the smoothness of differentiation.
4. The C4-RDE is a stable SDE.
5. Therefore, the observing eyes of the scientist and the doctor are considered to be the RDE, which consist of primitive-operators, or the primitive-operators themselves.

In the future, the SOC will show us not only inflammation, but elementary particle, the life and the new figure of the space–time continuum. Additional details have been provided below.

The diagnosis of the inflammation by the SOC is equivalent to the diagnosis of a new OOC from the SOC. However, the SOC already had the result when the sample was obtained. Accordingly, except for the manufacturing time, the calculation time of SOC is zero. If the calculation time is zero, the computer has P = NP property (Cook, 2000). Moreover, the relationship between gsOOC(s) and SOC is P = NP (Figure S7). A (conventional) computer does not include the manufacturing time in the calculation time.

The LCs (StrLC, SLL and MLC) described above accurately present cell and tissue injuries. Space information is represented by an empty space in a healthy tissue. The invasion is greater if the empty space is larger (even if the extent of cell injury is the same).

The analytical methods used in modern pathology are systems, in which cells are individually analysed. These approaches, including FCM, destroy tissue injury information. Transferring these concepts to inflammation assessment requires the measurement of leukocyte cluster information before the FCM because the FCM destroys the cluster of inflammatory cells. Importantly, the Lcnt destroys a cluster similarly.

WTLs and WSTLs combine when in close proximity, and a larger leukocyte cluster arises from a larger injury (be careful of the amount of supply of leukocytes). In contrast, a large cluster does not derive from a small injury. Therefore, the size of the leukocyte cluster (in a time zone) is proportional to the volume of the injury (in a time zone). This allows use of a staining solution to enable the observation of the changes in the injury in a time section and permits the representation of the injury as an SτC LC.

The ‘EOM’, including the NSE, can be easily applied in practice. However, such application of this equation is associated with several problems from AO or RO. These problems can be solved using the primitive-operator.

The SLL is a subset of the MLC. In the future, studies will be needed to analyse the inner time for the SLL (to MLC). The MLC can be used as a biomarker for C3-RDE (and C4-RDE). These RDEs are the SOC.

The volume of the injury at a certain time is proportional to the volume of the LC of the same *τ* cluster. Therefore, the LC can be used to calculate the velocity of inflammation and the antigen–antibody reaction. The RDE can operate the future, the past and the present of the LC.

Conventional mathematics, including the CDE, expresses only the world of S_2_. Therefore, a function can be used only in the first quadrant, and the problem changes to one associated with singularity and non-smoothness. If a function differentiates in the conventional-CS, in which S_1_ does not exist, a quadrant change is unnatural and not smooth. Str is eternal even if the function differentiates at any time. If the function does not use an Str operator (View operator), it cannot obtain smoothness.

The CDE is presented as follows:

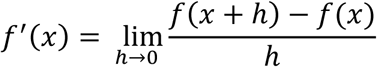

An example: *f*(*x*) = *ax*^2^ in the _R_Str (S_1_) side (Zai has inner Kyoku. S_1_ or S_2_ has no inner Kyoku.).

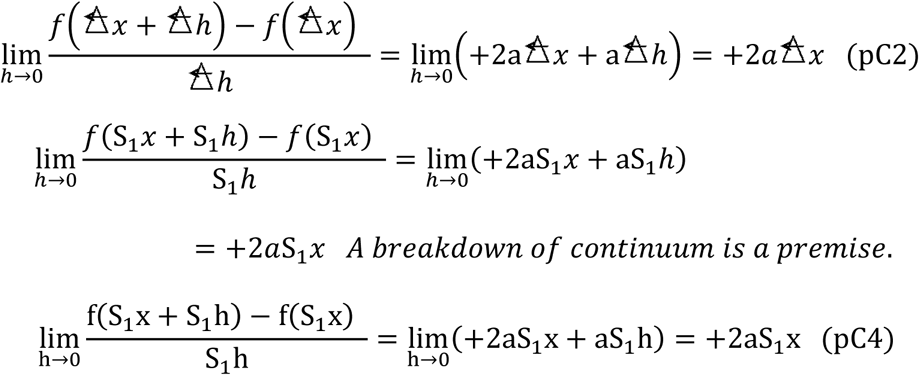

The limiting value in the CDE belongs in pC4 (Imaginary operator and value), which is a scale. Zai (value) (pC2) is described by an italic character. Kyoku (value) (pC1), pC3, pC5 and pC4 are described by standard characters. The correspondence to pC2 (real) from pC4 (imaginary) gives Zai. Therefore, StrLC is a transient form from CDE to RDE.

The numerator is bundled as follows:

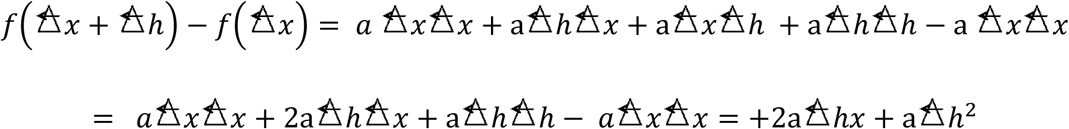

The denominator is bundled as follows:

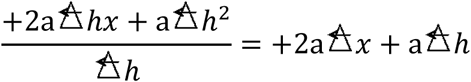

The relationship between the CS (StrZai) and the function is exact.

Str (S_2_) side:

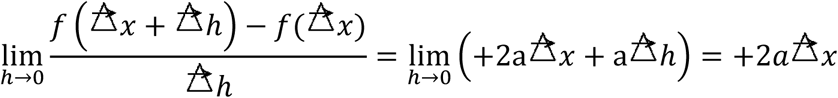

Exponent:

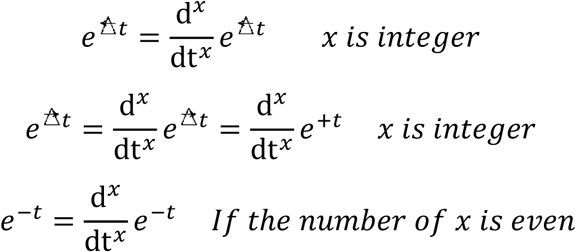

The CDE cannot determine Kyoku; therefore, the singularity and the integration constant are generated, allowing the convergence-differentiation using only one side of the CS.

If a conventional CS is established on an imaginary scale {imaginary-coordinate-system [ICS] in pC4 (Definition S1.5)}, the function involves a factor that represents a view (particularly Gv_1_), with the tool used to solve it as the Str operator. The view will change if a function mixing ‘–’ as the view (value) differentiates conventionally. This is unnatural and considered as the cause of the smoothness problem associated with differentiation. We must not mix absolute minus and relative minus without distinguishing them (i.e. relative operations _R_StrZai and _L_StrZai). The RCS for natural science should be formed from StrZai (Figure 6), with this view representing an observation. Therefore, the CS is the Zai (existence) itself and the observation. (IDP-EOM is improved by the C3-CS. A different-object interference is improved by the C4-CS. The NSE is improved by the C3-CS and/or C4-CS.)

The global view in a conventional-CS has to be distinguished from the function by the Str operator (particularly Gv_1_= –). The view is borrowed from the CS; hence, it must not be changed by conventional convergence-differentiation and a conventional absolute-value-operator. The conventional differential equation expresses only one side (_L_Str area), and the smoothness of a differential equation requires an Str operator similar to an EOM. The NSE is not equipped with Str operators and not smooth under all circumstances. The perfect answer to the Clay Millennium Problem (Fefferman et al., 2000) using the NSE is presented as follows:

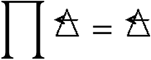

The limitations of the SOC, RDE, primitive-operators and SτC (LC) are currently unknown. Hence, research in this field should be continued to determine the limitations.

## ACKNOWLEDGEMENTS

I would like to thank the staff at Nonomura Dental Clinic for their clinical assistance; Dr. Sachiko Nisida and Dr. Hiroko Kamide for their assistance with language editing; Miss Haruko Kobayashi, Miss Manami Niwa, Miss Miki Matsushita, and Mr. Keita Taki for reference searches; and Mrs. Tomomi Okuda and Miss Saori Nishida for their assistance with the photographs and figures.

## Microscopes

The cover glass, slide glass, phase-contrast microscope and fluorescence microscope used herein were general models (Corporation Riken, Japan).

