## Supplementary for "Measurement and self-operating computer of the leukocyte continuum as a fixed space–time continuum in inflammation"

### Supplementary Texts

#### Supplementary Text S1

#### Background to the work as smoothness

From Leibniz [1], including La Geometrie of Rene Descartes [2] as orthogonal coordinate system, the solution of the equation of motion has been Figure S9. However, the real phenomenon is Figure S10 {as the orbiter of a motion object (of a simple substance)}. The term of acceleration is inconsistent. {It is a science fiction and comedy} Currently, applied mathematician and physicist shelve this problem and applied mathematics and physics are formed. From origin of the problem, through Absolute Origin (AO) and Relative Origin (RO), the article "**Fiendish million-dollar proof eludes mathematicians**" [3] is published. It has been simply misunderstanding from Leibniz [1] except Newton [4], at this paper publishing. This problem has erased the future of inflammation. In equation of motion with acceleration, when the direction which an observer sees is different, the result line is inconsistency about AO and RO. As the result of the "inconsistency line", modern science has various problems (caused by the imaginary coordinate system being based on imaginary mathematics): these problems are the directional-properties inconsistency problem of the equation of motion (Figure S9), our inability to solve the Navier–Stokes Equation [5, 6, and 3], the singularity (E.-W. Saw. et. al, 2016), the non-smoothness [5], the inconsistency of the integration constant, observation problem of quantum, and the instability of solutions to simultaneous differential equations [7-11] in applied mathematics and physics. However, only one Operator (StrZai) solves them. StrZai of the operator can also constitute Real Coordinate System (RCS) which indicates real space of Natural World.

#### Supplementary Text S2 Clinicians want to know tissue injury which becomes a basis of the diagnostic.

The diagnostic name of inflammation is named by giving '-itis' to name of living body tissue which has tissue injury. For example, periodontal tissue (attachment) is Periodontitis. In the case of the Periodontitis, if periodontal attachment (tissue) has tissue injury of loss (deepening) (dimension is L) and leukocyte infiltration (of mainly inflammatory cell) is observed to the tissue injury, it can do definitive diagnosis with Periodontitis [12]. However, clinically, dentists are doing the simple and rough diagnostics by deepening which is the past trace [13]. The form which indicates the extent of the tissue injury of periodontitis is measured by deepening (loss) by the one-dimensional approximation by probing. (Exactly, the form of $L^3T^{-1}$ is the extent of tissue injury.) Conventionally, in a certain inflammation, the form (Exactly, dimensions is $L^3T^{-1}$) of the injury tissue which indicates the extent of this injury and infiltration of leukocytes (Observing with the number with no dimension) were observed by the pathological tissue section [14]. (Tissue section observation has destruction of nonbonding and/or the inflammation tissue of weak coupling by surgical-removal extraction and section-izing. It means loss of an inflammation tissue to diagnose. It means that it is not necessary to predict future inflammation.)

Conventionally, secondary various biomarkers ($PGE_2$, Lactoferrin, Alkali phosphatase, Interleukin-1-β, Matrix metalloproteinase, Albumin, Aspartate aminotransferase, Lactate dehydrogenase, Secretor IgA) which are not essence were tried to inflammation, in view of the definition of inflammation [15]. Since they are not expressing the essence which is the above-mentioned definition, they have been stopped in a partial expression to diagnostics of inflammation and to measurement of the extent. As a result, in present clinic, the leukocyte count of the venous blood cannot do specification tissue of inflammation [16] or periodontitis have stagnated by low resolution form measurement to the tissue injury, such as the above, probing, and like roentgenography (CT and MRI etc. are included). They were far from the definitive diagnosis of inflammation and the quantification. Therefore, inflammation detects more than degree above the middle in the extent of disease, in the case of periodontitis, surgical treatment became a subject, and in cancer it was fatal.

**1** Leukocyte count problem

The leukocyte count from venous blood was proportion for an inflammation in living body [16]. In the local leukocyte count, in the clinic, the extent of inflammation by the local leukocyte count and periodontitis got the completely incoherent value, not to mention a proportional, like [17]. Since this phenomenon that occurs by the leukocyte count which is a primary biomarker, has been also generated by the secondary biomarker theoretically, completion of the primary biomarker by leukocytes is pressing need.

**2** The form measurement problem of tissue injury

The tomogram machine in cell level which can apply a form measurement of tissue injury to living body does not exist. Furthermore, in present-day pathology, the cell injury is the base [18], and measurement of the extent (form) of tissue injury is difficulty and low accuracy. (The above probing is the one example.) Even when the extent of cell injury are the same, the extent of tissue injury are greatly different exists without limit. Although the secondary biomarker generate from cell (injury), target of injury is the tissue injury which is space in a time division (interval).

**3** Simultaneous measurement problems {form (extent) of tissue injury & infiltration leukocytes}

Even if the above-mentioned trial is separately successful, there are simultaneous measurement problems of form (extent) of tissue injury and the infiltration leukocytes.

In the conventional leukocyte count method [16], it was destroying this space information (exactly time-space information) and was mixing (= destroying) multi tissue injury information.

Of course, flow cytometry is also the same. If the time- division (interval) is not distinguished, the size of injury becomes unknown. It is understood by leukocyte cluster with time division (interval). Especially In the injury of a periodontal pocket, when it passes through an injury part, it must be able to observe and calculate the time division (interval) of LC(Leukocyte continuum). By statistics sampling method (includes leukocyte count and flow cytometry [19] etc.) [16 and 17], it was destroying this time-space information (in leukocyte and tissue injury).

Accordingly this problem that was not able to measure an extent of inflammation {the number of inflammatory cell (mainly leukocyte) infiltration and measurement of the form which is the extent of injury} was not a biological problem but a measurement science (physics) problem.

The problems of diagnostics (arithmetic value in the future, the present, and the past by the number of constructed leukocytes and the form) was not completed by the measurement value were also physical and an applied mathematics problem.

In the time-space information of (detection of) tissue injury and leukocyte infiltration, It is necessary to insert the relationship of tissue injury (time-space problem) and cell injury (biological problem) at the concept of pathology, clearly.    Supplementary

**Supplementary Text S3**

History of inflammation

The present inflammation measurement can measure the extent of an inflammation with the leukocyte count of venous blood. However, an important diagnostic name like the above-mentioned is not detected.

And the Virchow era shows that the true character of inflammation is leukocyte infiltration [14], and thing [14] which inflammation generates according to tissue injury, accordingly a tissue name with injury constitutes a diagnostic name of inflammation,

As a result, invasive sick extent must have been able to measure according to the inflammation which is all the invasive sick responses (defense reaction of a living body). (Leukocyte infiltration by the microscope observation under the tissue injury of the 'burn' of the tongue of the frog by Cohnheim [14], and tongue tissue)

However, in present condition, Cautions tend only toward the injury of a cell, important tissue injury is driven away out of the system of pathology [14 and 18].

Like the reference 6, the extent of periodontitis and leukocyte count gets a completely incoherent value, it is not a proportional relationship.

However, according to experimental research, the number of leucocytes and periodontitis also has proportional data, the extent and the leukocyte count of periodontitis have a paradox problem.

From two above-mentioned reports, the cause of the problem was not able to extract a sample from a simplex tissue injury part, but a sample will take synthetic sampling from the few complex tissue injury parts (multi tissue injury) (with micro orders).

Including periodontal pocket, currently, the micro sampling which positions a simplex tissue injury part with a micro level is impossible. Furthermore, the tomogram with 1-micron level which can sample leukocytes is also next to impossible in clinical for human.

**Supplementary Text S4**

Secondary biomarkers

$PGE_2$, Alkaline phosphatase, Interleukin-1 beta, Matrix metalloproteinase, Lactoferrin, Albumin, Aspartate aminotransferase, Lactate dehydrogenase, Secretory IgA, They are generated from a cell in the tissue injury and the other tissue injury.

**Supplementary Text S5**

**No ethics problems**

I have posted using clinical data for journal, and such as society, on the noticeboard of my clinic (Figure S0). Therefore, there is no ethics problem.

**Supplementary Text S6**

**6.1** $S_1$ side is (Minus side), (Figures 6 and S10)  it is solution below, by integration

$$m\frac{d^2y}{dt^2} = \left(t\triangle\right)^0 ma = \triangle ma \text{ 、 } \frac{d^2y}{dt^2} = \triangle a$$

In this case, m is mass. Other is m is phase value.

$$\int \frac{d^2y}{dt^2}dt = v = \frac{dy}{dt} = a\left(\mathrm{t}\triangle\right)^1 + C_0\left(\mathrm{t}\triangle\right)^0 = a\left(\mathrm{t}\triangle\right)^1 + v_0\triangle$$

$$\int \frac{dy}{dt}dt = y = \frac{1}{2}a\left(\mathrm{t}\triangle\right)^2 + C_0\left(\mathrm{t}\triangle\right)^1 + C_1\left(\mathrm{t}\triangle\right)^0 = \frac{1}{2}a\left(\mathrm{t}\triangle\right)^2 + v_0\left(\mathrm{t}\triangle\right)^1 + y_0\triangle$$

$$y = \frac{1}{2}a\left(\mathrm{t}\triangle\right)^2 + v_0\left(\mathrm{t}\triangle\right)^1 + y_0\triangle$$

If it differentiates here

$$\frac{dy}{dt} = \triangle a\mathrm{t}^1 + \triangle v_0 \;, \qquad \frac{d^2y}{dt^2} = \triangle a$$

{Str (**Definition S2**)is constant on an independent axis (G axis), and Potential is set to $t^2$ or $t^1$. t is physical time. The Physical time t (Figure S13) constitutes a Zai from the uncertainty principle. Conventionally, the time has been reported as an object like a bubble.}

Accordingly, The Real Coordinate System (RCS) is formed as StrZai($\triangle$and$\triangle$), and time ( $t^1$ , $t^2$, and $t^q$ …) which is the Po is given. (In each term, refer to Figure 6**.** And the solutions are y = $y_1+y_2$ include C3 Coordinate system, Figure legend 6.)

**6.2** $S_2$ side is (Plus side) (Figures 6 and S10**)** (Example S2)

**6.3** Definition Space is real space of Natural World.
RCS indicate real space of Natural World. Because, It element-izes our Natural World exactly. (Table S1, Definition S1)

**Supplementary Text S7**

**Background to the work as SDE**

Since Natural World is space time continuum (in brain time space continuum), the analysis of phenomenon in science depends on the Differential Equation. Furthermore, In order to solve the phenomenon in which two or more kinds interfere, Simultaneous Differential Equation (SDE) is needed. However, the SDE cannot solve in many cases [7-11]. The problem is coordinate system problems (Especially smoothness problem), lack of Str (View) Operator **(Definition S2)**, lack of Relative Differentiation Equations (RDE), The RDE built to base on the Primitive Operator which element-ized Nature by phase classification. And, Since the RDE has Real Coordinate System as StrZai, it can solve the problem. Therefore, this paper rebuilt conventional SDE by the simultaneous RDE as an example the equation of Prey & Predator. So, It obtained a stable solution.

**Supplementary Materials**

**Supplementary Material S1**

The conventional equation of motion,

$$m\frac{d^2y}{dt^2} = \pm ma$$

(a: acceleration, m:mass in the Material S1, y:distance, t:time) (Example S1)
The solution is (Figure S9) in the mathematics after Leibniz [1]. This differs from the 'right line' in the Principia [4] which Newton says. Furthermore, the solution differs from the actual phenomenon (Figure S10). (The origin in this case is an Absolute Origin. At a RO, the 1st quadrant also produces this inconsistent phenomenon. It is also a smoothness problem of NSE. Furthermore, this problem is involved also in generating of singularity deeply.)

**1.1** The conventional relative equation of motion
Even if the equation (1.1) extends to the relative equation, the relative equation (with ICS) has the same inconsistency. The equation (1.1) is a fundamental example. You can change equation (1.1) into relative equation.
The acceleration and the velocity are parallel relationship.

$$F = m\gamma\frac{dv}{dt} + m\gamma^3\left(\frac{v^2}{c^2}\cdot\frac{dv}{dt}\right)$$

In this case, $\gamma = \left(1 - {v^2}/{c^2}\right)^{-\frac{1}{2}}$ , $\frac{dv}{dt} = a = \frac{d^2y}{dt^2}$ , $v\ is\ velocity\ on\ y\ axis$

The acceleration and the velocity are orthogonal relationship.

$$\boldsymbol{F} = m\gamma \frac{dv}{dt} + m\gamma^3 \left( \frac{vu}{c^2} \cdot \frac{du}{dt} \right) \quad , \quad u \; is \; velocity \; on \; x \; axis$$

**1.2** Other equation of motion

The other equation (with ICS) has the same inconsistency.

$$\frac{D^2\boldsymbol{s}}{D\tau^2} = \frac{D\boldsymbol{V}}{D\tau} = \mathbf{a} \quad , \qquad \mathbf{a} = (\mathrm{a}_x, \mathrm{a}_y, \mathrm{a}_z), \qquad \boldsymbol{s} = (x, y, z), \qquad \boldsymbol{V} = (u, v, w)$$

An example is Navier-Stokes's Equation (except compression term)

$$\mathrm{a}_x = \frac{D^2x}{D\tau^2} = \frac{Du}{D\tau} = \frac{\partial u}{\partial \tau} + u\frac{\partial u}{\partial x} + v\frac{\partial u}{\partial y} + w\frac{\partial u}{\partial z} = -\frac{1}{\rho}\frac{\partial P}{\partial x} + \nu\nabla^2 u + F_x$$

$$a = \mathrm{a}_y = \frac{D^2y}{D\tau^2} = \frac{Dv}{D\tau} = \frac{\partial v}{\partial \tau} + u\frac{\partial v}{\partial x} + v\frac{\partial v}{\partial y} + w\frac{\partial v}{\partial z} = -\frac{1}{\rho}\frac{\partial P}{\partial y} + \nu\nabla^2 + F_y$$

$$\mathrm{a}_z = \frac{D^2z}{D\tau^2} = \frac{Dw}{D\tau} = \frac{\partial w}{\partial \tau} + u\frac{\partial w}{\partial x} + v\frac{\partial w}{\partial y} + w\frac{\partial w}{\partial z} = -\frac{1}{\rho}\frac{\partial P}{\partial z} + \nu\nabla^2 + F_z$$

$$\nabla^2 = \frac{\partial^2}{\partial x^2} + \frac{\partial^2}{\partial y^2} + \frac{\partial^2}{\partial z^2}$$

(a: acceleration, *x*, *y*, *z* : positions on the coordinate, *u*, *v*, *w :* velocity on *x*, *y*, *z*, F : external force, p pressure, ρ density, νcoefficient of kinematic viscosity)

**Supplementary Material S2**

**Supplementary Definition S1**

**1.1 ω1_Operator** for the acquisition of primitive operator

Nature World (NW) is classified according to phase classification, by ω1_Operator.

**1.1.1** ω1_Operator is, (pF is the symbol of Phase Field. Phase Field is integer field.)

$$World = {}^{\omega d}\begin{bmatrix} \mathrm{Ex} \\ \mathrm{Np} \\ \mathrm{Wp} \end{bmatrix}_{\mathrm{pF}} = \begin{bmatrix} \mathrm{Ex} \\ \mathrm{Np} \\ \mathrm{Wp} \end{bmatrix}_{\mathrm{pF}} = \begin{bmatrix} \mathrm{Ex} \\ \mathrm{Np} \\ \mathrm{Wp} \end{bmatrix}, \qquad \begin{matrix} Ex = World.Ex = World \\ Np = World.Np \\ Wp = World.Wp \end{matrix}$$

ω1_Operator classifies Ex according to Np and Wp in right-hand side which are properties (ω character value), and generates each world in left-hand side. Observer exists on an axis. (It's named G axis.). Fundamentally, Observer exists on G axis. (Be careful of an observer's position.)

**1.1.2** Ex

Ex expresses the value of the Operator. Ex is a number or a function. Value of Ex is integer or real number.

**1.1.3** ω Character value : Np and Wp

**1.1.3.1** Np (the Number by phase)

Np indicates countable thing by phase. (Np is the number of thing by phase.) It is integer.

The Number by phases (Np) is number of Kyoku counted by the phase on G axis.

**1.1.3.2** Wp (the Width by phase)

Wp indicates the thing which has width by phase. (Wp is the width of thing by phase.) It is integer. If existence is more than two, they distinguish by Po. (Po is real number or integer.) The Width by phase (Wp) is the width between Kyoku and Kyoku which were contrasted by the phase on G axis. (Distance which can be indicated by the phase)

**1.1.3.3** Example of Np and Wp for an Object,

Example of Np for an Object,

$\bar{0}$ : expresses the number which is not zero. It is countable Object

0 : Zero,

Example of Wp for an Object,

$\bar{0}$ : Object with width,

0 : Object without width,

**1.2** Phase class 1 (pC1);

$$| = {}^{\omega}\begin{bmatrix} \mathrm{Ex} \\ 0 \\ 0 \end{bmatrix}_{\mathrm{n}} \qquad (1.1)$$

is Kyoku |, (The Ex is a orthogonal relation to G axis) shows the boundary between existence and existence, or the boundary between existence and space-time. Kyoku constitutes K axis.

**1.3** Phase class 2 (pC2);

$$🜁 \text{ or } 🜁 \text{ or } 🜁 = {}^{\omega}\begin{bmatrix} \mathrm{Ex} \\ \bar{0} \\ \bar{0} \end{bmatrix}_{\Delta_n} \qquad (1.2)$$

is Zai 🜁, 🜁, and 🜁 which is constituted by closed Kyoku. It is inner Kyoku (iK). (Ex is a real parallel relation to G axis. $\bar{0}$ expresses the number which is not 0.) Zai shows the existence in Nature World (NW). Zai constitutes G axis. Zai is Closed Field. It is Inner Field. On the other hand, Single Kyoku makes the Open Field.

**1.4** Phase class 3 (pC3);

$$\Sigma| = {}^{\omega}\begin{bmatrix} \mathrm{Ex} \\ \bar{0} \\ 0 \end{bmatrix}_{\mathrm{n}} \qquad (1.3)$$

is Set of Kyoku, Σ|, (Ex is a real and/or Image orthogonal relation to G axis)

**1.5** Phase class 4 (pC4);

$$\leftarrow \text{ or } \rightarrow = {}^{\omega}\begin{bmatrix} \mathrm{Ex} \\ 0 \\ \bar{0} \end{bmatrix}_{\mathrm{n}} \qquad (1.4)$$

is Image Interval (Ex is an image parallel relation to G axis ) This is a vector.

**1.6** Phase

After all, the phase is the world divided (partitioned) by the Kyoku. The phases are the contrast (comparison) divided (partitioned) by the Kyoku.

**1.6.1** phase time and phase space

Examples are, phase time is a time which is divided (partitioned) by the Kyoku, and phase space is a space which is divided (partitioned) by the Kyoku.

**1.7** Pure Number (pN)

pN is out of the classification, it is class5 (pC5). Pure number doesn't have Np and Wp. The element number of Np and Wp, it is pEN. In Np, It is one piece in Nature World even if it divides into a half. The unit of 0.5 pieces does not exist actually in Nature World.

In example of this paper, n, m, and k are phase value as arbitrary positive integers. Here, phase is the world divided by the Kyoku.

**1.8** Real Mathematics and Imaginary Mathematics

By ω1_Operator, Real Mathematics is pC1, pC2 and the part of pC3. Imaginary Mathematics is pC4 and the part of pC3.

**2** Kyoku and Zai are primitive operator.

**Supplementary Definition S2 (Basic)**

View and Str, (primitive operator)

View in the same area is constant.

$$\mathrm{View} = \mathrm{View} * \mathrm{View} \quad (2.1)$$

$$\mathrm{Str} = \mathrm{View} * \mathrm{pEN} \quad (2.2)$$

$$\mathrm{Str} = \mathrm{Str} \cdot \mathrm{Str} \quad (2.3)$$

pEN =1, pEN is pure Element Number. '*' is Constant Multiplication. Even if Constant Multiplication multiplies the same View, it is the constant multiplication character. View in the same area is constant. The kind of View is Local View, Global View, and Relative View.

The kind of sign value in View: Lv=+ (Only), $Gv_1$= –, $Gv_2$=+, $Rv_1$= –, $Rv_2$ = +

$$\mathrm{Gv}_1 = \mathrm{Gv}_1 * \mathrm{Gv}_1, \ \ \mathrm{Gv}_2 = \mathrm{Gv}_2 * \mathrm{Gv}_2 \quad (2.4)$$

$$\{\mathrm{Rv}_1 \text{ or } \mathrm{Rv}_2\} = (\mathrm{Gv}_1 \text{ or } \mathrm{Gv}_2) * \mathrm{Lv} \quad (2.5)$$

$${}_{\mathrm{R}}\mathrm{Str} = \mathrm{Rv}_1 * \mathrm{pEN} = \mathrm{S}_1, \quad {}_{\mathrm{L}}\mathrm{Str} = \mathrm{Rv}_2 * \mathrm{pEN} = \mathrm{S}_2$$

The conventional operator (especially '–') is absolute operator. However, Str operators (${}_{R}$Str and ${}_{L}$Str) are relative operator. ${}_{R}$StrZai and ${}_{L}$StrZai have different View. Conventional minus (-) operator is absolute operator.

**Supplementary Definition S3 (Basic)**

**Primitive Operators (Closed Field in a part of Primitive Operator) (Basic)**

**3.1** UZai (Primitive Operator is element Operator.)

pC2 of ω1_Operator generated the Uncertainty Zai (UZai) as the simple element in Nature World (NW). .

$$\Delta = \begin{bmatrix} ? \\ ? \\ ? \end{bmatrix}_{\Delta_n} \qquad (3.1)$$

The { or } is Uncertain Operator. ? = {–1 or +1}

**3.2** The right Str Zai $\left({}_{R}\mathrm{StrZai}\right)$(Secondary Operator) is,

$${}_{n}\triangle_{n+m} = {}_{n}\triangle_{n+1} = {}^{\omega}\begin{bmatrix} S_1 \cdot g(m) \\ S_1 \cdot g(m) \\ S_1 \cdot g(m) \end{bmatrix}_{{}_{n}\triangle_{n+1}} = g(m) \cdot \begin{bmatrix} S_1 \\ S_1 \\ S_1 \end{bmatrix}_{\triangle} = S_1 \cdot \begin{bmatrix} g(m) \\ g(m) \\ g(m) \end{bmatrix}_{\triangle} = \triangle$$

$$= -1 \quad (3.2)$$

Fundamentally, m = 1, $g$(m)=1, $g_2$(m) = n – (n–m) = +m, or $g_1$(m) = (n+m) –n = +m, from Local Field. See Definition S3. (Basic)

Our world is Relative Field. In Relative Field, right Str Zai is –1. and In Local Field, value of the right Str Zai is +1. The difference in the value is a difference in an observer's position.

**3.3** The left Str Zai $\left({}_{L}\mathrm{StrZai}\right)$ (Secondary Operator) is,

$${}_{n-m}\triangle_{n} = {}_{n-1}\triangle_{n} = {}^{\omega}\begin{bmatrix} S_2 \cdot g(m) \\ S_2 \cdot g(m) \\ S_2 \cdot g(m) \end{bmatrix}_{{}_{n-1}\triangle_{n}} = g(m) \cdot \begin{bmatrix} S_2 \\ S_2 \\ S_2 \end{bmatrix}_{\triangle} = S_2 \cdot \begin{bmatrix} g(m) \\ g(m) \\ g(m) \end{bmatrix}_{\triangle} = \triangle$$

$$= +1 \quad (3.3)$$

**3.4** Secondary Operator

Secondary Operator consists of two or more element operators. As an example, Str Zai consists of View and UZai. (UZai is quantum element is elementary particle)

**3.5** The template of Str, $S_?$

$S_?$Zai is Macro Uncertainty Zai (MuZai) (Secondary Operator)

$${}_{n-m}\triangle_{n} \text{ or } {}_{n}\triangle_{n+m} = \begin{bmatrix} S_? \cdot m \\ S_? \cdot m \\ S_? \cdot m \end{bmatrix}_{\triangle_n} = \triangle \cdot m \quad (3.4)$$

$$S_? = \left\{ {}_{R}\mathrm{Str} \text{ or } {}_{L}\mathrm{Str} \right\} = \{ S_1 \text{ or } S_2 \}$$

($S_?$ which is the template of Str does not change by differentiation. However, the '?' is changed by differentiation.) (The real example is single leukocyte.)

**3.6** Relative Coordinate System (Real Coordinate System) (RCS) △△ △△

RCS consists of △ and/or △, which is/are real. RCS consists of the scale by IDO {Inner Field Draws (Defines) Outer Field} and an origin by IOSF (Inner Kyoku-and-Outer Kyoku Synchronous Field) (iK and oK).

**3.6.1** C3 Relative Coordinate System is indicated by a phase in detail, (m=1, Po=1), (3.5)
This example is the phase n which has origin by IOSF, and has the scale (n-m, n+m) by IDO. Fundamentally, m = 1.

| | | | | | |
|---|---|---|---|---|---|
| | △Po | | △Po | | $= {}_{n-m}\Delta_{n}\ {}_{n}\Delta_{n+m}$ |
| $+m$ | $+m$ | $0\ 0$ | $+m$ | $+m$ | Local Field $\{Lv * pEN \cdot g_3(m)\}$ |
| $n-m$ | $S_1 m$ | $n\ n$ | $S_2 m$ | $n+m$ | Relative Field $(Gv * \text{Local Field})$ |

| | | | | | |
|---|---|---|---|---|---|
| | △Po | | △Po | | $= {}_{n-m}\Delta_{n}\ {}_{n}\Delta_{n+m}$ |
| $+m$ | $+m$ | $0\ 0$ | $+m$ | $+m$ | Local Field $\{Lv * pEN \cdot g_3(m)\}$ |
| $n-m$ | $- * m$ | $n\ n$ | $+ * m$ | $n+m$ | Relative Field $(Gv * \text{Local Field})$ |

Phase value $g_3(m)$ & $g_3(m)$ Local Field

$g_3(m) = (n+m) - n = +m$

$g_{31}(m)$ & $g_{32}(m)$ Relative Field

$g_{31}(m) = (n+S_1m) - n = + S_1m = -m$ $g_{32}(m) = (n+S_2m) - n = S_2m = +m$

**3.6.2** C4 Relative Coordinate System, (3.6)

| | | | | | |
|---|---|---|---|---|---|
| | △Po | | △Po | | $= {}_{n-m}\Delta_{n}\ {}_{n}\Delta_{n+m}$ |
| $-m$ | $+m$ | $0\ 0$ | $+m$ | $-m$ | Local Field $\{Lv * pEN \cdot g_4(m)\}$ |
| $n-m$ | $S_2 m$ | $n\ n$ | $S_1 m$ | $n+m$ | Relative Field $(Gv * \text{Local Field})$ |

| | | | | | |
|---|---|---|---|---|---|
| | △Po | | △Po | | $= {}_{n-m}\Delta_{n}\ {}_{n}\Delta_{n+m}$ |
| $-m$ | $+m$ | $0\ 0$ | $+m$ | $-m$ | Local Field $\{Lv * pEN \cdot g_4(m)\}$ |
| $n-m$ | $+ * m$ | $n\ n$ | $- * m$ | $n+m$ | Relative Field $(Gv * \text{Local Field})$ |

Phase value $g_4(m)$ & $g_4(m)$ Local Field

$g_4(m) = n - (n-m) = +m$

$g_{41}(m)$ & $g_{42}(m)$ Relative Field

$g_{41}(m) = n - (n - S_2m) = +m$ $g_{42}(m) = n - (n - S_1m) = S_1m = -m$

Left hand side in the equation: The number just under △ is the value of StrZai in the each field, the both sides is phase values, the upper row is a phase of Local Field, the lower row is Relative

Field, and the row is a phase of the Inner and Outer Complex Field. Right hand side in the equation indicates the detail of Relative Field notation. {In this case, $g$(m)=1} Our world is Relative Field, in Relative Field, $\triangle$= –1. In Local Field, $\triangle$= +1. The difference in the value is a difference in an observer's position.

**Supplementary Notation S1**

Abbreviated notation

C3 is $^{R}(^{L}\triangle)\,^{R}(^{L}\triangle)$, C4 is $^{R}(^{L}\triangle)\,^{R}(^{L}\triangle)$. More, C3 is $\triangle\triangledown\triangle$, C4 is $\triangle\triangledown\triangle$.

**1.1** Multi Scale

Zai continuum is Multi Scale by IDO

Example of $\triangle\triangledown\triangle\triangledown\triangle$ are n-3, n-2, n-1, n, and $\triangle\triangledown\triangle\triangledown\triangle$ are n, n-1, n+1, n+2, n+3.

**1.2** The same kind of StrZai are operated at the Local Field, finally, different StrZai are operated by Relative Field.

**Supplementary Definition S4** Smooth-Qualitiy

**4.1** The Definition of Smooth-Quality

The value of Parallel Four Arithmetic Operations (PFAO) always is constant in all area? The value or sign of each operation always is the value of (Table S1) in all area?

**4.2** Smooth-Quality of conventional ICS in pC4, and RCS in pC2, pC3 and pC1

Result of conventional ICS and RCS in Smooth-Quality is shown (Table S1). The RCS have Smooth-Quality in the all Parallel Four Arithmetic Operations (PFAO). However, the ICS does not have Smooth-Quality in Parallel Multiplication (PM) and Parallel Division (PD). It is Rough-Quality. Therefore, in the ICS, the multiplication and division operate correctly only at part of the $1^{st}$ quadrant by Absolute Origin. (Be careful Relative Origin in $1^{st}$ quadrant) Furthermore, in the ICS, the result shows that there is no smoothness in the conventional differentiation (CD). This is also answer of a Millennium Problem NSE. [5]

**4.3** Continuum PM examines Conventional Coordinate System (ICS) and RCS $\triangle$ (continuum)

**4.3.1** ICS in pC4 is, (→is the plus vector)

$$\prod_{m=0}^{k} {}_{n+m}\rightarrow_{n+(m+1)} = {}_{n}\rightarrow_{n+1} = +1$$

**4.3.2** RCS in pC2 is (The origin n do not influence the operation.)

$$\prod_{m=0}^{k} \overset{\text{Local or Relative}}{{}_{n+m}\triangle_{n+(m+1)}} = \overset{\text{Local or Relative}}{{}_{n}\triangle_{n+1}} = {}^{\text{Rf or Lf}}\triangle = {}^{\text{Lf}}\triangle = {}^{\text{Rf}}\triangle = +1$$

Local and Global View of $_{L}$Str

$$_{n}\dashrightarrow = +$$

+ area (on independent variable axis) of the Relative or Local Coordinate System is Smooth-Quality in PM.

**4.4 C**ontinuum PM examines Conventional Coordinate System (ICS)

In minus area(− or $S_1$ area) of conventional ICS in pC4, at least one of all the n and k is,

(n, n≧0, m, and k are k>m≧0 in arbitrary integers) (←is the minus vector)

$$_{n-1}\leftarrow_{n} = \leftarrow = -1$$

$$\prod_{m=0}^{k} {}_{n+(-m-1)}\leftarrow_{n-m}$$

The m is even number, the infinite product value is -1. The m is odd number, the value is +1. In – area (minus sign area) of the conventional ICS in pC4, Smooth-Quality is not filled. It is Rough-Quality. Furthermore, in + area, multiplication of the minus vector takes Rough-Quality.

**Supplementary Definition S5** Bundling

(n, m and k are k>m≧0 in arbitrary integers,) Observer position is in Real Field (RF).

**5.1** Continuum Zai (Phase Locking type) bundle

$_{R}$StrZai (x is arbitrary integers)

$$\prod_{j=0}^{x} {}_{n+1}\overleftarrow{\triangle}_{n} = \overleftarrow{\triangle}^{x} = \overleftarrow{\triangle} = -1$$
$$n-1 \quad n$$

$_{L}$StrZai

$$\prod_{j=0}^{x} {}_{n}\overrightarrow{\triangle}_{n+1} = \overrightarrow{\triangle}^{x} = \overrightarrow{\triangle} = +1$$
$$n \quad n+1$$

Abbreviated,

$$\overleftarrow{\triangle} = \overleftarrow{\triangle}^{x}$$

It describes only the $_{R}$Str, $_{L}$Str is also similar. Continuum StrZais is calculated as a single StrZai. (*x* is arbitrary integers)

**5.2** $_{R}$StrZai $\overleftarrow{\triangle}$ (continuum) (Phase Shift type) is,

$$\prod_{m=0}^{k} \underset{n+(-m-1)}{{}_{n+(m+1)}}\triangle_{\underset{n+(-m)}{n+m}} = \underset{n-1}{{}_{n+1}}\triangle_{\underset{n}{n}} = \triangle = -1$$

**5.3** Abbreviated, (phase shift type and phase locking type)

$$\prod \triangle = \triangle$$

This is defined as Bundling. And, this equation is smooth.

**5.4** Global View in $_{R}$Str,

$$\dashleftarrow_{n} = -$$

An infinite division is the same. Other parallel 3 arithmetic operations have no problem like the above. Therefore, C3 and C4 coordinate system in pC2 are Smooth-Quality.

**5.5** Bundling and *Distributing* in Po

$$\triangle \cdot \underset{\leftarrow}{{}_{n+S_1 m}t_n} \;\; \underset{Po\ Distributing}{\overset{\text{Po Bundling}}{\underset{\leftarrow}{\overset{\rightarrow}{=}}}} \;\; \triangle \cdot t$$

t is pure number or Kyoku number, $t$ is Zai number.

**5.6** *Bundling* and *Distributing* in Zai

$$\prod \triangle \;\; \underset{Zai\ Distributing}{\overset{Zai\ Bundling}{\underset{\leftarrow}{\overset{\rightarrow}{=}}}} \;\; \triangle$$

**Supplementary Notation S2** Local-Relative Operation (LRO) $^{R}(^{L}\triangle)$, $^{R}(^{L}\triangle)$

Operation of Local Field to Relative Field (LRO)

Therefore, PM in a RCS is,

$$^{R}(^{L}\triangle) \cdot {}^{R}(^{L}\triangle) = {}^{R}(^{L}\triangle) = {}^{R}\triangle = {}^{R}\triangle \cdot {}^{R}\triangle = \triangle \;, \quad \triangle = {}^{R}(^{L}\triangle) = {}^{R}(^{L}\triangle \cdot {}^{L}\triangle)$$

$$^{R}(^{L}\triangle) \cdot {}^{R}(^{L}\triangle) = {}^{R}(^{L}\triangle) = {}^{R}\triangle = {}^{R}\triangle \cdot {}^{R}\triangle = \triangle \;, \quad \triangle = {}^{R}(^{L}\triangle) = {}^{R}(^{L}\triangle \cdot {}^{L}\triangle)$$

These operators harmonize with the Natural World. Furthermore,

$$^{R}\triangle = {}^{R}(^{L}\triangle) = {}^{R}(^{L}\triangle) = {}^{R}\triangle = \triangle \;, \quad {}^{R}\triangle = {}^{R}(^{L}\triangle) = {}^{R}\triangle = \triangle$$

**Supplementary Example S1 (an example of CDE)**

Equation of motion is solution (in time-space continuum: TSC) below, by integration (Cx, x is integration constant, is given a number with every from 0 in appearance order of integration constant)

$$m\frac{d^2y}{dt^2} = \pm ma \quad 、 \quad \frac{d^2y}{dt^2} = \pm a$$

$$\int \frac{d^2y}{dt^2}dt = \frac{dy}{dt} = \pm at^1 + C_0 = v = \pm at^1 + v_0$$

$$\int \frac{dy}{dt}dt = \pm\frac{1}{2}at^2 \pm C_0t^1 + C_1 = y = \pm\frac{1}{2}at^2 \pm v_0t^1 + y_0$$

The 2nd term of the right-hand side(RHS) indicates a smooth motion and smooth flow, however the 1st term of the RHS has indicated that presence of external force, such as a gravity, is acting on a motion object, a flow, and etc. {The 1st term of the RHS is a motion which rebounds a rigid body (wall) etc. The flow is not matching at the 2nd term of the RHS. They are unnatural.} The direction of a motion and flow is reversed at an Absolute or Relative Origin. Those origins were the conventional singularity.

**Supplementary Example S2** $S_2$ side is (Plus side)

$$\frac{d^2y}{dt^2} = \vec{\triangle} a$$

$$v = \vec{\triangle} at^1 + \vec{\triangle} v_0$$

$$y = \vec{\triangle}\frac{1}{2}at^2 + \vec{\triangle} v_0t^1 + \vec{\triangle} y_0$$

**Supplementary Example S3** If $f(x) = ax^3$ is differentiated,

$$\lim_{h\to 0}\frac{f\left(\vec{\triangle}x + \vec{\triangle}h\right) - f\left(\vec{\triangle}x\right)}{\vec{\triangle}h} = +3a\vec{\triangle}x\vec{\triangle}x = +3a\vec{\triangle}(x\cdot x) = +3a\vec{\triangle}x^2$$

Here, the denominator is,

$$a\left(\vec{\triangle}x + \vec{\triangle}h\right)^3 - a\left(\vec{\triangle}x\right)^3$$

$$= \left(a\vec{\triangle}x\vec{\triangle}x + a\vec{\triangle}h\vec{\triangle}x + a\vec{\triangle}x\vec{\triangle}h + a\vec{\triangle}h\vec{\triangle}h\right)\left(\vec{\triangle}x + \vec{\triangle}h\right) - a\vec{\triangle}x\vec{\triangle}x\vec{\triangle}x$$

$$= a\vec{\triangle}x\vec{\triangle}x\vec{\triangle}x + a\vec{\triangle}h\vec{\triangle}x\vec{\triangle}x + a\vec{\triangle}x\vec{\triangle}h\vec{\triangle}x + a\vec{\triangle}h\vec{\triangle}h\vec{\triangle}x$$
$$+a\vec{\triangle}x\vec{\triangle}x\vec{\triangle}h + a\vec{\triangle}h\vec{\triangle}x\vec{\triangle}h + a\vec{\triangle}x\vec{\triangle}h\vec{\triangle}h + a\vec{\triangle}h\vec{\triangle}h\vec{\triangle}h$$
$$-a\vec{\triangle}x\vec{\triangle}x\vec{\triangle}x$$

$$= +3a\vec{\triangle}h\vec{\triangle}x\vec{\triangle}x + 3a\vec{\triangle}h\vec{\triangle}h\vec{\triangle}x + a\vec{\triangle}h\vec{\triangle}h\vec{\triangle}h$$

And,

$$\frac{+3a\triangle h\triangle x\triangle x+3a\triangle h\triangle h\triangle x+a\triangle h\triangle h\triangle h}{\triangle h}$$

$$=+3a\triangle\cdot\triangle x\triangle x+3a\triangle h\triangle\cdot\triangle x+a\triangle h\triangle h\triangle$$

$$=3a\triangle x^2+3a\triangle hx+a\triangle h^2 \underset{\leftarrow Distributing}{\overset{Bundling \rightarrow}{=}} \triangle(3ax^2+3ahx+ah^2)$$

**Supplementary Definition S6**

Relative Differential Equation Basic model

N is the number of observation object, Po is Potential, Po is τ time, τ is phase time (in this paper is biological time as phase.), the m is phase and n is phase origin. γ is the coefficient of solution. Pure Element Number is pEN=1. Outer Kyoku is,

$$\frac{\begin{array}{c}K\\ |\,\mathrm{N}\\ \tau_n\end{array}}{\begin{array}{c}Z\\ |\\ \tau_n\end{array}}=\frac{\begin{array}{c}K\\ |\,\mathrm{N}_\tau\\ n\end{array}}{\begin{array}{c}Z\\ |\,\tau\\ n\end{array}}=\frac{{}_r\mathrm{d}_{\tau_n}\mathrm{N}}{{}_r\mathrm{d}_{\tau_n}}=\frac{{}_r\mathrm{d}_n\mathrm{N}_\tau}{{}_r\mathrm{d}_n\tau}=\frac{{}_r\mathrm{dN}_{\tau_n}}{{}_r\mathrm{d}\tau_n}$$

**6.1 Various condition models** (SτC in SOC)

C is condition, the each condition as the following.

C1 and C2 from C3

$$\text{Condition 1(C1)(Existence 1):}\quad \frac{{}_r\mathrm{dN}_{\tau_n}}{{}_r\mathrm{d}\tau_n}=\frac{{}_{n-m}\triangle_{n}\,N_{2\tau}}{{}_{n-m}\triangle_{n}\,\tau}-\frac{{}_{n}\triangle_{n+m}N_{1\tau}}{{}_{n}\triangle_{n+m}\tau}$$

$$=+\frac{{}_{n+S_1m}\triangle_{n}\,N_{2\tau}}{{}_{n+S_1m}\triangle_{n}\,\tau}+\frac{{}_{n}\triangle_{n+S_2m}N_{1\tau}}{{}_{n}\triangle_{n+S_2m}\tau}=\frac{{}_{n+S_1m}\triangle_{n}\,N_{2\tau}}{{}_{n+S_1m}\triangle_{n}\,\tau}-\frac{{}_{n}\triangle_{n+S_2m}N_{1\tau}}{{}_{n}\triangle_{n+S_2m}\tau}$$

$$\text{Condition 2(C2)(Existence 2):}\quad \frac{{}_r\mathrm{dN}_{\tau_n}}{{}_r\mathrm{d}\tau_n}=-\frac{{}_{n-m}\triangle_{n}\,N_{2\tau}}{{}_{n-m}\triangle_{n}\,\tau}+\frac{{}_{n}\triangle_{n+m}N_{1\tau}}{{}_{n}\triangle_{n+m}\tau}$$

$$= \frac{{}_{n+S_1m}\triangle_{n}\, N_{2\tau}}{{}_{n+S_1m}\triangle_{n}\, \tau} + \frac{{}_{n}\triangle_{n+S_2m} N_{1\tau}}{{}_{n}\triangle_{n+S_2m} \tau} = -\frac{{}_{n+S_1m}\triangle_{n}\, N_{2\tau}}{{}_{n+S_1m}\triangle_{n}\, \tau} + \frac{{}_{n}\triangle_{n+S_2m} N_{1\tau}}{{}_{n}\triangle_{n+S_2m} \tau}$$

C3 and C4 use Relative Origin (Kyoku) (C3 is the most familiar.) (Figure S7)

Condition 3(C3): $\frac{{}_{r}d_{\tau_n}N}{{}_{r}d_{\tau_n}} = +\frac{{}_{n-m}\triangle_{n}\, N_{2\tau}}{{}_{n-m}\triangle_{n}\, \tau} + \frac{{}_{n}\triangle_{n+m} N_{1\tau}}{{}_{n}\triangle_{n+m} \tau} = +\frac{{}_{n-m}\triangle_{n}\, N_{L\tau}}{{}_{n-m}\triangle_{n}\, \tau} + \frac{{}_{n}\triangle_{n+m} N_{R\tau}}{{}_{n}\triangle_{n+m} \tau}$

$$= +\frac{{}_{n+S_1m}\triangle_{n}\, N_{2\tau}}{{}_{n+S_1m}\triangle_{n}\, \tau} + \frac{{}_{n}\triangle_{n+S_2m} N_{1\tau}}{{}_{n}\triangle_{n+S_2m} \tau}$$

Condition 4(C4): $\frac{{}_{r}dN_{\tau_n}}{{}_{r}d\tau_n} = +\frac{{}_{n-m}\triangle_{n}\, N_{2\tau}}{{}_{n-m}\triangle_{n}\, \tau} + \frac{{}_{n}\triangle_{n+m} N_{1\tau}}{{}_{n}\triangle_{n+m} \tau}$

$$= +\frac{{}_{n-S_2m}\triangle_{n}\;{}_{2}N_{\tau}}{{}_{n-S_2m}\triangle_{n}\, \tau} + \frac{{}_{n}\triangle_{n-S_1m}\;{}_{1}N_{\tau}}{{}_{n}\triangle_{n-S_1m} \tau}$$

C4 special notation of a different kind $\frac{{}_{r}dN_{\tau_n}}{{}_{r}d\tau_n} = +\frac{{}_{n-m}\triangle_{n}\;{}_{2}N_{\tau}}{{}_{n-m}\triangle_{n}\, \tau} + \frac{{}_{n}\triangle_{n+m}\;{}_{1}N_{\tau}}{{}_{n}\triangle_{n+m} \tau}$

Calculation of τ

$$\tau_{n+S_1m} = \tau \cdot (n + S_1m)\ , \qquad \tau_{n+S_2m} = \tau \cdot (n + S_2m),$$

Primitive condition is

Condition 0(C0)(primitive RDE): $| = {}_{n-?m}\Delta_n + {}_{n}\Delta_{n-?m}$

As for n and m, phase (value). The n indicates a Relative Origin (Kyoku |). t is t toki. τ is τtoki. indicates the time of τ. The n, m, and l indicate a phase (value) (n≧0,m≧0, l≧0). The variable may use arbitrary variable.

**6.2** γ (for OOC)

C1, C2, C3 is,

$$\frac{{}_{\mathrm{r}}\mathrm{dN}_{\tau_n}}{{}_{\mathrm{r}}\mathrm{d}\tau_n} = \frac{{}_{\mathrm{r}}\mathrm{d}_{\tau_n}\mathrm{N}}{{}_{\mathrm{r}}\mathrm{d}_{\tau_n}} = \quad \gamma\,\mathrm{N} = \left( -\frac{{}_{\mathrm{n-m}}\overset{\rightarrow}{\triangle}_{\mathrm{n}}\, N_{2_\tau}}{{}_{\mathrm{n-m}}\overset{\rightarrow}{\triangle}_{\mathrm{n}}\, \tau} - \frac{{}_{\mathrm{n}}\overset{\leftarrow}{\triangle}_{\mathrm{n+m}} N_{1_\tau}}{{}_{\mathrm{n}}\overset{\leftarrow}{\triangle}_{\mathrm{n+m}} \tau} \right)$$

$$\gamma = \left( -\frac{{}_{\mathrm{n-m}}\overset{\rightarrow}{\triangle}_{\mathrm{n}}\, N_{2_\tau}}{{}_{\mathrm{n-m}}\overset{\rightarrow}{\triangle}_{\mathrm{n}}\, \tau} - \frac{{}_{\mathrm{n}}\overset{\leftarrow}{\triangle}_{\mathrm{n+m}} N_{1_\tau}}{{}_{\mathrm{n}}\overset{\leftarrow}{\triangle}_{\mathrm{n+m}} \tau} \right) \frac{1}{\mathrm{N}} = |\gamma$$

N is, (The following | | is RAO.) (The condition 4 has a reverse sign)

$$\mathrm{N} = \left( \left| -{}_{\mathrm{n-m}}\overset{\rightarrow}{\triangle}_{\mathrm{n}}\, N_{2_\tau} \right| + \left| -\,{}_{\mathrm{n}}\overset{\leftarrow}{\triangle}_{\mathrm{n+m}} N_{1_\tau} \right| \right) \cdot \frac{1}{2}$$

Therefor

$$\gamma = \left( \frac{-{}_{\mathrm{n-m}}\overset{\rightarrow}{\triangle}_{\mathrm{n}}\, N_{2_\tau} - {}_{\mathrm{n}}\overset{\leftarrow}{\triangle}_{\mathrm{n+m}} N_{1_\tau}}{\left| -{}_{\mathrm{n-m}}\overset{\rightarrow}{\triangle}_{\mathrm{n}}\, N_{2_\tau} \right| + \left| -\,{}_{\mathrm{n}}\overset{\leftarrow}{\triangle}_{\mathrm{n+m}} N_{1_\tau} \right|} \right) \frac{2}{\tau} = |\gamma$$

C4 is

$$\gamma = \left( \frac{{}_{\mathrm{n-m}}\overset{\rightarrow}{\triangle}_{\mathrm{n}}\, N_{2_\tau}}{{}_{\mathrm{n-m}}\overset{\rightarrow}{\triangle}_{\mathrm{n}}\, \tau} + \frac{{}_{\mathrm{n}}\overset{\leftarrow}{\triangle}_{\mathrm{n+m}} N_{1_\tau}}{{}_{\mathrm{n}}\overset{\leftarrow}{\triangle}_{\mathrm{n+m}} \tau} \right) \frac{1}{\mathrm{N}} = |\gamma$$

$$\mathrm{N} = \left( \left| {}_{\mathrm{n-m}}\overset{\rightarrow}{\triangle}_{\mathrm{n}}\, N_{2_\tau} \right| + \left| {}_{\mathrm{n}}\overset{\leftarrow}{\triangle}_{\mathrm{n+m}} N_{1_\tau} \right| + \right) \cdot \frac{1}{2}$$

Therefor

$$\gamma = \left( {}_{\mathrm{n-m}}\overset{\rightarrow}{\triangle}_{\mathrm{n}}\, N_{2_\tau} + {}_{\mathrm{n}}\overset{\leftarrow}{\triangle}_{\mathrm{n+m}} N_{1_\tau} \right) \frac{1}{N \cdot \tau} = |\gamma,$$

$$\gamma = \left( \frac{{}_{\mathrm{n-m}}\overrightarrow{\triangle}_{\mathrm{n}}\, N_{2_{\tau}}}{\left|{}_{\mathrm{n-m}}\overrightarrow{\triangle}_{\mathrm{n}} N_{2_{\tau}}\right|} + \frac{{}_{\mathrm{n}}\overleftarrow{\triangle}_{\mathrm{n+m}} N_{1_{\tau}}}{\left|{}_{\mathrm{n}}\overleftarrow{\triangle}_{\mathrm{n+m}} N_{1_{\tau}}\right|} \right) \frac{2}{\tau} = |\gamma,$$

**6.3** OOC example of the solution and τSC

A certain solution (variable separation, target function etc.) inserts above-mentioned γ. (γ is Kyoku)

If an initial value is, $e^{\gamma t_n}$, n is Origin , m is nonOrigin,

$$\tau_{n+m} = (n + m) \cdot \tau$$

$$\mathrm{Str}_1 \cdot \tau_{n+m} = \mathrm{Str}_1 \cdot (n + m) \cdot \tau = (n + \mathrm{Str}_1 \cdot m) \cdot \tau$$

$$\mathrm{Str}_2 \cdot \tau_{n+m} = \mathrm{Str}_2 \cdot (n + m) \cdot \tau = (n + \mathrm{Str}_2 \cdot m) \cdot \tau$$

In decreasing function,

$$N_{\tau_{n+m}} = e^{\gamma(t_n + \mathrm{Str}_1 \cdot \tau_{n+m})}$$

In increasing function,

$$N_{\tau_{n+m}} = e^{\gamma(t_n + \mathrm{Str}_2 \cdot \tau_{n+m})}$$

They are smooth.

**6.4** AO and RO

The ‘n’ is Origin. It is AO or RO. The phase ‘n’ which $_{\mathrm{R}}$StrZai and $_{\mathrm{L}}$StrZai meet or leave is ‘AO’. The conjugation Kyoku of Zais of the different Str is ‘AO’. The conjugation Kyoku of Zais of the same Str is ‘RO’. AO and RO of LF are the same.

**Supplementary Method S1**

**1** Method of using StrZai

In order to solve this problem, use Str Operator defined below.

$$\prod \overleftarrow{\triangle} = \overleftarrow{\triangle} \quad 、 \quad \prod \overrightarrow{\triangle} = \overrightarrow{\triangle}$$

$$| = \overrightarrow{\triangle} + \overleftarrow{\triangle} = 0 , \qquad | = \overleftarrow{\triangle} + \overrightarrow{\triangle} = 0$$

$\overleftarrow{\triangle}$ and $\overrightarrow{\triangle}$ are StrZai which are Uncertainty Zai with Str (Operator). | is Kyoku. ‘1’ is pure number. The details of StrZai, Zai, Kyoku and pure number are indicated by Definitions S1, S2, S3, S4, S5 and another paper (detail) “Re-calculation of modern civilization by the New-Operator which Phase Class makes”.

$\overleftarrow{\triangle}$\{$_{\mathrm{R}}$Str ($S_1$)\} or $\overrightarrow{\triangle}$\{ $_{\mathrm{L}}$Str ($S_2$)\} is defines as Local Field. Both Local Fields always have a value of + for the Zai inner value (a local independent variable). View of a Local Field is also only +. When two or more Local Fields are combined, each View changes relatively. View is an observer's observation direction. Therefore, View should not be mixed with a function. Both should be discriminated.

Two combining fields are defined as Relative Field. This relative field is formed as C3 coordinate system. (Figure 6) In the case of the motion simple substance in this paper, ${}_{R}$Str ($S_1$) side is as minas side (View is minas - ) and ${}_{L}$Str ($S_2$) side is as plus side (View is plus + ).

(At the combining field in the antigen-antibody reaction of leucocytes and microorganisms, this thing is understood easily. The combining field is C4.)

I apply Str Operator to the conventional differentiation, and solve the previous problem. Concretely, I set an independent variable axis as Single StrZai (or StrZai continuum).

In actual usage, If real number or distinction is required, the Zai is multiplied by ‘Po’. As an example, it is ‘t’. (In this case’t’ is time.) (Figure 6)

**1.1** Operation of ωd (switch) of ω1_Operator

$$\mathrm{Po}\ {}^{ON}\begin{bmatrix}\mathrm{Ex}\\\mathrm{Np}\\\mathrm{Wp}\end{bmatrix}_{\mathrm{pF}} = \begin{bmatrix}\mathrm{Po\ Ex}\\\mathrm{Np}\\\mathrm{Po\ Wp}\end{bmatrix} = \begin{bmatrix}\mathrm{Ex\ Po}\\\mathrm{Np}\\\mathrm{Wp\ Po}\end{bmatrix},\ {}^{ON}\begin{bmatrix}\mathrm{Ex}\\\mathrm{Np}\\\mathrm{Wp}\end{bmatrix}_{\mathrm{pF}}\ \mathrm{Po} = \begin{bmatrix}\mathrm{Po\ Ex}\\\mathrm{Po\ Np}\\\mathrm{Po\ Wp}\end{bmatrix} = \begin{bmatrix}\mathrm{Ex\ Po}\\\mathrm{Np\ Po}\\\mathrm{Wp\ Po}\end{bmatrix}$$

If this ωd (switch) is ON, the ω1_Operator can distinguish the width and the number.

$$\mathrm{Po}\ {}^{OFF}\begin{bmatrix}\mathrm{Ex}\\\mathrm{Np}\\\mathrm{Wp}\end{bmatrix}_{\mathrm{pF}} = {}^{OFF}\begin{bmatrix}\mathrm{Po\ Ex}\\\mathrm{Po\ Np}\\\mathrm{Po\ Wp}\end{bmatrix}_{\mathrm{pF}} = {}^{OFF}\begin{bmatrix}\mathrm{Ex\ Po}\\\mathrm{Np\ Po}\\\mathrm{Wp\ Po}\end{bmatrix}_{\mathrm{pF}} = {}^{OFF}\begin{bmatrix}\mathrm{Ex}\\\mathrm{Np}\\\mathrm{Wp}\end{bmatrix}_{\mathrm{pF}}\ \mathrm{Po}$$

If this ωd (switch) is OFF, the ω1_Operator have commutative law.

**Supplementary Material S3** Solution of Simultaneous Differential Equation

Condition 4 (C4) Relative Interference Differential Equation

$$\frac{\substack{\mathrm{K}\\ |\mathrm{N}_{\tau}\\ \mathrm{n}}}{\substack{\mathrm{K}\\ |\tau\\ \mathrm{n}}} = \frac{\substack{\mathrm{K}\\ |\ \mathrm{N}\\ \tau_{\mathrm{n}}}}{\substack{\mathrm{K}\\ |\\ \tau_{\mathrm{n}}}} = \frac{{}_{\mathrm{r}}|\mathrm{N}_{\tau_{\mathrm{n}}}}{{}_{\mathrm{r}}|\tau_{\mathrm{n}}} = \frac{{}_{\mathrm{r}}\mathrm{dN}_{\tau_{\mathrm{n}}}}{{}_{\mathrm{r}}\mathrm{d}\tau_{\mathrm{n}}} = \frac{{}_{\mathrm{r}}\mathrm{d_n N}_{\tau}}{{}_{\mathrm{r}}\mathrm{d_n}\tau} = \frac{\triangle_{\mathrm{n}\ 2}N_{\tau}}{\triangle_{\mathrm{n}}\tau} + \frac{\triangle_{\mathrm{n}\ 1}N_{\tau}}{\triangle_{\mathrm{n}}\tau}$$

r is an Identifier of Relative Differentiation. n and m are phase. (integer) τ is τ_time (potential).

N is the number (positive integer). ${}_{1}$N and ${}_{2}$N are the existence 1 and the existence 2. $\triangle$ is ${}_{L}$StrZai, $\triangle$ is ${}_{R}$StrZai. Str means Stream in inertia system and Strepto in static system.

**Supplementary Definition S7**

**S7.1** Definition of Active leukocyte (AL), destructive leukocyte after W(S)TL

It corresponds to biological time as phase time by Kyoku change. (Convertible biological time and physical time)

**7.1.1** Classification by inner Kyoku function (**Definition S1**) (Active function or Destructive of function)

Active leukocyte has inner Kyoku (iK) function. In this case, iK is membrane. Primary membrane is cell membrane. Secondary membrane is nuclear membrane.

Destructive leukocyte doesn't have inner Kyoku (iK) function.

**7.1.2** Classification by the presence or absence of inner Kyoku (iK)

Presence iK leukocyte has inner Kyoku. Leukocyte with iK

Absence iK leukocyte doesn't have inner Kyoku. Leukocyte without iK

**7.1.3** Example of AL, DL and DLfc

**7.1.3.1** Staining

Detection of the inner Kyoku function by staining, and the further visualization of inner Kyoku,

Detection of the boundary function (activity or in-activity) by staining, (Determination of the degree of activity by staining,)

The classification of the degree of activity by acid red (FigureS8)

Active leukocyte: AL0, AL1, AL2, (Short-distance combining leukocytes)

Destructive leukocyte: DL0, DL1, DL2 (Dissociation of leukocytes)

AL0: no depletion

→AL1 Nuclear membrane depletion, Nuclear Inner Kyoku depletion,

→AL2 Cell membrane depletion, Cell Inner Kyoku depletion,

→DL0 (Both) Inner Kyoku functional stop (Dissociation of leukocytes)

→DL1 Membrane breakdown, Inner Kyoku of cell break-down, (There is cell membrane)

AL0 to DL1 is presence iK and DL2 is absence iK

→DL2 Cell membrane extinction, Inner Kyoku extinction of cell, (The cell membrane nothing, only nuclear)

→DLfc Generating of fibrous connective tissue, (Addition of fibrous tissue)

AL and DL corresponds biological time. Example ALn is $\tau_n$ (Figure S8). Example of biological is '$\tau$=1'.

Correspondence of biological time and physical time is future work.

**7.1.3.2** No staining

**7.1.3.2**.1 Classification by function

Active leukocyte: There is inner Kyoku function. (The function by inner Kyoku is kept up) There is motion of a granule.

Destructive leukocyte: There is no inner Kyoku function. (The function by inner Kyoku is not kept up) There is no motion of a granule.

**7.1.3.2.2** Classification by presence iK or absence iK

Active leukocyte: There is inner Kyoku. There is cell membrane.

Destructive leukocyte: There is not inner Kyoku. There is no cell membrane.

**S7.2** Definition of SτC and τSC

From above-mention (Definition S6.1 and S6.3),

**7.2.1** The contour of Leukocyte itself expresses space.

**7.2.2** ALs combines together in short distance. (Figure 1) As a result, ALs forms LC.

**7.2.3** AL and DL expresses biological time as phase time.

Therefore, the cluster by W(S)TL forms the biological (fixed) space-time continuum.

This time is the time relatively fixed to each leukocyte. (And, the change of Kyoku is the phase change. Accordingly, the change of Kyoku time is the phase time change.) It is simultaneous time. The fixed (simultaneous) time is τ time. The τ is phase time. Here, phase is the world divided by the Kyoku (Definition S1). I define this continuum as SτC in SOC and as τSC in OOC.

**7.2.4** An example is

$$\frac{{}_{\mathrm{n+S_1 m}}\triangle_{\mathrm{n}}\, N_{2_\tau}}{{}_{\mathrm{n+S_1 m}}\triangle_{\mathrm{n}}\, \tau} \quad \text{(SOC) is S}\tau\text{C.}$$

$${}_2\mathrm{N}_{\tau_\mathrm{n}} = {}_2\mathrm{N}_0 \cdot \exp\left({}_2\gamma_{\tau_\mathrm{n}} \cdot \tau_\mathrm{n}\right), \quad \mathrm{N} = f(\tau) \quad \text{(OOC) is }\tau\text{SC.}$$

W(S)TL has inner field. The inner field separate Outer Kyoku (space). Therefore, The inner field has inner Kyoku as envelope. (Figure S4, S5, S6, and S7)

**7.2.5**

$$\frac{\Delta space}{\Delta \tau} \quad \text{(SOC) is S}\tau\text{C.} \qquad\qquad \text{Space} = f(\tau) \quad \text{(OOC) is }\tau\text{SC.}$$

SτC is space fixed-time continuum. τSC is fixed-time-space continuum. A solution of SOC is OOC.

$$\frac{space}{\Delta\tau}\ or\ \frac{S_? \cdot space}{\Delta\tau} : \ space\ is\ without\ iK, it\ is\ no\ continuum, \text{(it is S}\tau\text{I).}$$

SτI is space fixed-time intermittent. The space is intermittent. It cannot be bundled. However, if a trace is predicted, it can be bundled. (STI is space-time intermittent.) (Time has iK.)

### Supplementary references

### Mathematical and physical references


### Biological references

### Supplementary Tables

| | Conventional ICS | | RCS | |
|---|---|---|---|---|
| | $S_1$ area (−area) | $S_2$ area(+area) | $S_1$ area (−area) | $S_2$ area(+area) |
| Para. Multi. | +1 = −1·(−1) | +1 = +1·(+1) | −1 = △ = △ · △ | +1 = △ = △ · △ |
| Para. Div. | +1 = −1÷(−1) | +1 = +1÷(+1) | −1 = △ = △ ÷ △ | +1 = △ = △ ÷ △ |
| Para. Add | −2 =−1+(−1) | +2 = +1+(+1) | −2 = △ + △ | +2 = △ + △ |
| Para. Sub. | 0 =\|=−1−(−1) | 0 =\|= +1−(+1) | 0 = \| = △ − △ | 0 = \| = △ − △ |

**Table S1 Survey of Smooth-Quality conventional ICS and RCS**

Conventional Imaginary Coordinate System (ICS) use absolute minus. Real Coordinate System (RCS) consists of pC2 and pC1 (+ partially pC3). (It use relative minus) The ICS does not have Smooth-Quality. However, the RCS have Smooth-Quality. Kyoku is generated by Parallel Subtraction (Para. Sub.), in Zai Pair.

| inside | Area 1 | Area 2 | Area 3 | Area 4 | outside |
|---|---|---|---|---|---|
| Area | 438313 | 424228 | 414979 | 524449 | |
| γ as Area | 0.0326593 | 0.0220422 | -0.233057 | | |
| N | 476 | 286 | | | |
| γ as N | 0.4986877 | | | | |

**Table S2 By OOC, Area, N and γ in MLC (the value in LF)**

Detailed OOC is future work, Here Please see SOC! The inflammation which was being cured has become acute. This table calculated Figure S6. General drawing software (MICROGRAFX DRAW. 7) and general Picture processing software (EasyAccess_67123) was used.

| No | Size of dot image (H*V) | H(μm) | V(μm) |
|---|---|---|---|
| 1 | 99*93 | 17.028 | 17.4375 |
| 2 | 93*86 | 15.996 | 16.1250 |
| 3 | 90*85 | 15.480 | 15.9375 |
| 4 | 81*77 | 13.932 | 15.1875 |
| 5 | 92*82 | 15.824 | 15.3750 |

| | | | |
|---|---|---|---|
| 6 | 88*81 | 15.136 | 15.1875 |
| 7 | 81*70 | 13.932 | 13.1250 |
| 8 | 76*70 | 13.072 | 13.1250 |
| 9 | 85*72 | 14.620 | 13.5000 |
| 10 | 84*75 | 14.448 | 14.0625 |
| 11 | 89*82 | 15.308 | 15.3750 |
| 12 | 98*87 | 16.856 | 16.3125 |
| 13 | 84*87 | 14.448 | 16.3125 |
| 14 | 89*83 | 15.308 | 15.5625 |
| 15 | 91*75 | 15.652 | 14.0625 |
| | mean | 15.136 | 15.1125 |
| | deviation | 1.088 | 1.283 |

**Table S3**

**Size of WTL**

| File | Condition of Upper Leukocyte and time | Condition of Lower Leukocyte and time |
|---|---|---|
| BMP450.bmp | AL0 1:40 | AL0 1:40 |
| BMP959.bmp | AL0 2:06 | AL0 2:06 |
| BMP1160.bmp | AL1 4:05 | AL0 4:05 |
| BMP1583.bmp | AL1 4:24 | AL1s 4:24 |
| BMP1817.bmp | AL2s 4:46 | AL1 4:46 |
| BMP1988.bmp | AL2 4:52 | AL2s 4:52 |
| BMP2297.bmp | AL2 5:27 | AL2 5:27 |

**Table S4 Condition of StrLC**

**s is start of stage.**

### Supplementary Figures legend

**Figure S1 Multi Circle Explorer (MCE)**

The scale is 1mm. This **MCE** is manufactured by Isizuka Co., Ltd. (Japan)

411-3 Kiurishinden, Yoshikawa-city, Saitama, Japan 342-0044

In this paper, the cover grass, the slide glass, the phase-contrast microscope, and the fluorescence microscope is general model. (Manufactured by MicroDent)

**Phase-contrast microscope** and **fluorescence microscope** manufactured by MicroDent.

Mail:

Limited company MicroDent

**Figure S2 Detail measurement line of Figure 1**

The contact line (center Red line is contact line in Figure 1) of the Leukocyte 1 (L1) and the Leukocyte 2 (L2) is drawn. It is Bond length (Bl). Two straight lines (side Red lines are Leukocyte L1 and Leukocyte L2 in Figure 1) are drawn on the diameter parallel to the contact line and widest. The blue straight line {is Dist (Distance line) in Figure 1} is drawn on a contact line and a right angle so that it pass through the center of a contact line. (Scale 10 µm)

**Figure S3 The leukocytes in blood do not combine, versus Figure 2**

(A): Time 0:00:00 is start. (B): Time 0:00:30 is two leukocytes meet and contact. (C): Time 0:00:56 is two leukocytes still contact. (D): Time 0:01: 04 is two leukocytes leave. (Scale 10 µm)

**Figure S4 The proportional size of the injury predicted by SLL of Figure 2**

White envelope line: The envelope of Leucocyte Cluster of the same τ by dynamic movements of cell organelles is proportional to the size of injury. RO (Relative Origin) and AO (Absolut Origin) is conventional singularity. The Leukocyte boundary has ability of RO or AO. (see Figure S7) (Scale is same as Figure 2)

**Figure S5 MLC is SτC indicate space and time, as detail of Figure 5A, refer to Figure S6,**

(A, C and E): fluorescence microscope image, (B and D): Phase contrast microscope image, (E): An example of τ boundary and τ zone, the arrow indicates a space series and time series. And, the apex (it is AO) indicates the singularity to the future of inflammation. (F): An example of StrLC, Scale of (A) and (E): 100 µm, (B) ,(C) and (D): 20µm, (F): 10µm

**Figure S6 For SOC and OOC, τ boundary and τ zone as Figure 5A LC**

(A): is the figure (B) made to overlap a real image as Figure S5 (A to E) LC. (B): An example of τ boundary and τ zone. This LC is same as Figure S5 (A to E) LC. Since this inflammation is macroscopic, the microscope image is patched. (It is a limit of an optical system.) And, CNL of the image painted by red. (LC is Leukocyte continuum.)

**Figure S7 The RDE as LC, for SOC and OOC in Figure 5A, S5, S6, and 7,**

The ‘n’ is Origin. The ‘n’ of end of the arrow of time series is AO. The phase ‘n’ which $_{R}$StrZai and $_{L}$StrZai meet or leave is ‘AO’. The conjugation Kyoku of Zais of the different Str is ‘AO’. The conjugation Kyoku of Zais of the same Str is ‘RO’. AO and RO of LF are the same.
(A) Example of MuZai and UZai, (B) $_{R}$StrZai and $_{L}$StrZai as C1 RDE. (C) C1 RDE as SτC. The end of the arrow of a time series is ‘AO’. (see Figures S5 and S6) (D) C4 RDE by microorganism and leukocyte. (In A, B, and D, A part of Po abbreviates. )

**Figure S8 Conventional equation of motion**

A conventional solution of equation of motion in conventional coordinate system from Leibniz, (It has “inconsistency line” which is red line in second quadrant.)

**Figure S9 Equation of motion with Str Operator (C3 coordinate system)**

An explanation of each line is, (orthogonal axis is y, and lateral is StrZai t.)

$$\text{red line: } \frac{1}{2}a\left(t\overset{\nwarrow}{\triangle}\right)^{2}\ \{{}_{\mathrm{R}}\text{Str side}(\mathrm{S}_1\text{ side})\},\quad \frac{1}{2}a\left(t\overset{\nearrow}{\triangle}\right)^{2}\ \{{}_{\mathrm{L}}\text{Str side}(\mathrm{S}_2\text{ side})\}\ \ \text{(C3)},$$

$$\text{blue line: } v_0\left(t\overset{\nwarrow}{\triangle}\right)^{1}\{{}_{\mathrm{R}}\text{Str side}(\mathrm{S}_1\text{ side})\},\qquad v_0\left(t\overset{\nearrow}{\triangle}\right)^{1}\ \{{}_{\mathrm{L}}\text{Str side}(\mathrm{S}_2\text{ side})\}\ \ \text{(C3)},$$

$$y_0\left(t\overset{\nwarrow}{\triangle}\right)^{0}+y_0\left(t\overset{\nearrow}{\triangle}\right)^{0}=0,\qquad y_0\overset{\nwarrow}{\triangle}+y_0\overset{\nearrow}{\triangle}=0=|\,y_0\ \ at\text{ origin}\ \ \text{(C3)}$$

$$\frac{1}{2}a\left(t\overset{\nwarrow}{\triangle}\right)^{2}+\frac{1}{2}a\left(t\overset{\nearrow}{\triangle}\right)^{2}=0,\quad v_0\left(t\overset{\nwarrow}{\triangle}\right)^{1}+v_0\left(t\overset{\nearrow}{\triangle}\right)^{1}=0\qquad \text{(C3)}$$

$$\mathrm{t}\ \geq 0$$

The singularity is clear! C3 coordinate system solves the problem of the singularity, the integration constant and the initial value. The red line is ‘right line’. Newton said ‘right line’ in LAW2 of Principia. {(Isacc Newton 1687) [4]} LAW2 of the book said, “The alteration of motion is ever proportional to the motive force impressed; and is made in the direction of the right line in which that force is impressed. “ Newton and Leibniz is branch point in history.

**Figure S10 Example of C4 Relative Differentiation and C4 RCS as Figure 7**

In the conventional equation, leukocytes or microorganisms were calculated as antimatter. Exactly, View (Operator) was mixing to the function. Even if it views the right and views the left in the space (coordinate system), cell does positive proliferation (multiplication). (Scale 10 μm)

**Figure S11 Example of C4 Relative Differentiation and C4 RCS as Figure 7**

(Scale 10 μm)

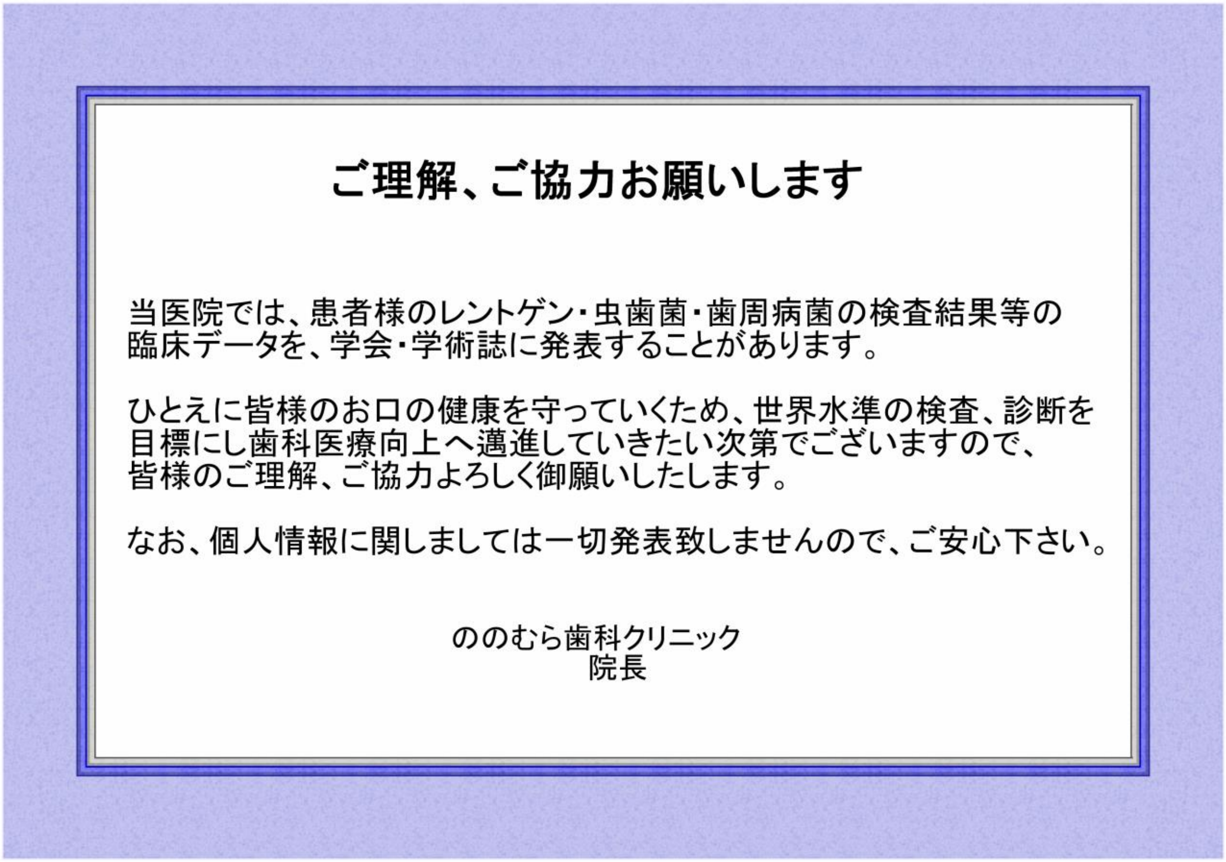


**Figure S0 no ethics problems**

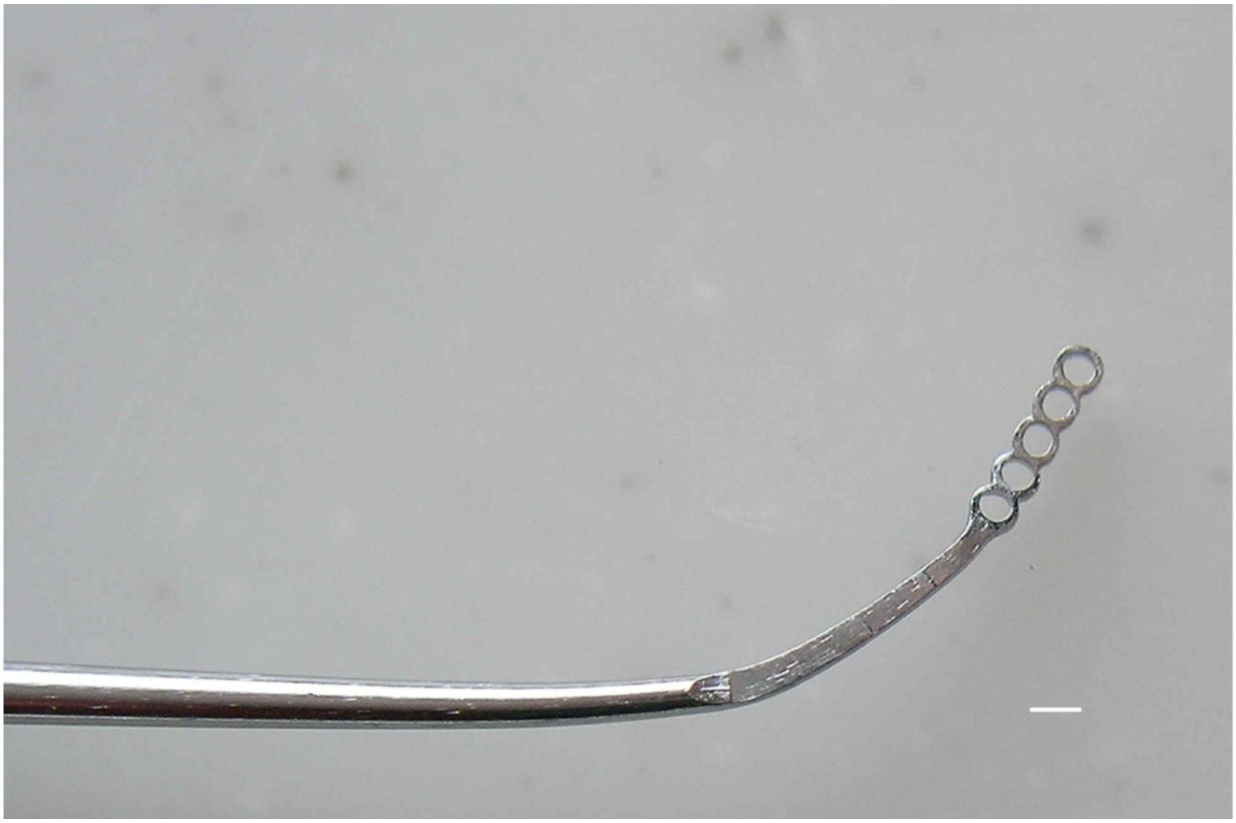

**Figure S1 Multi Circle Explorer (MCE)**

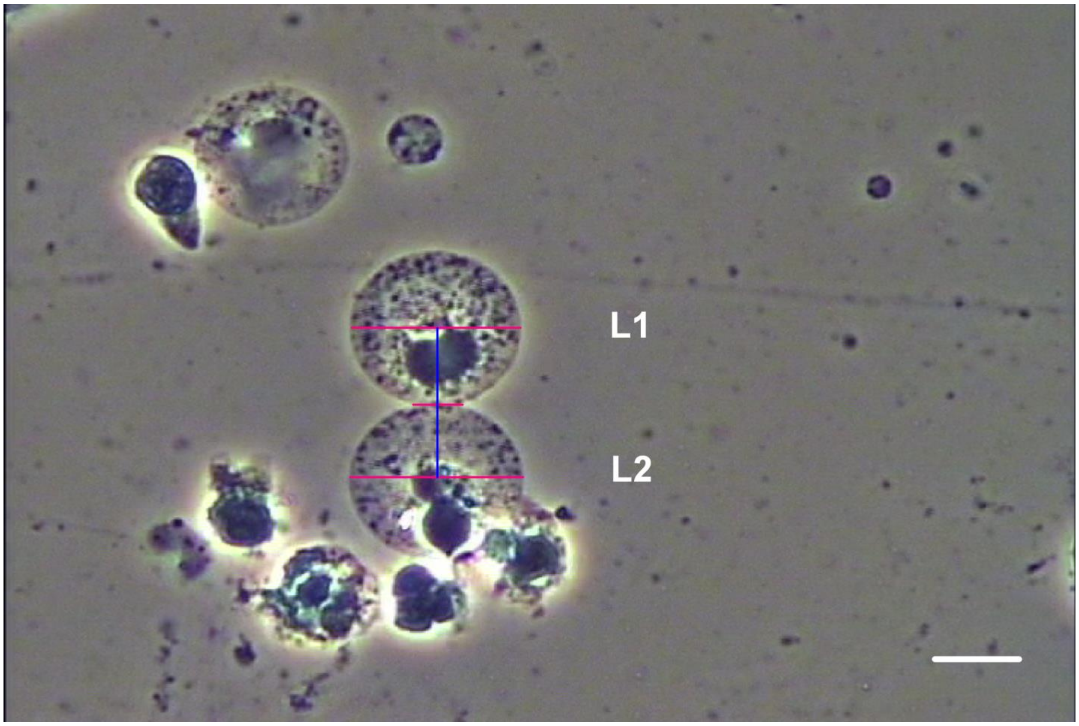


**Figure S2 Detail measurement line of Figure 1**

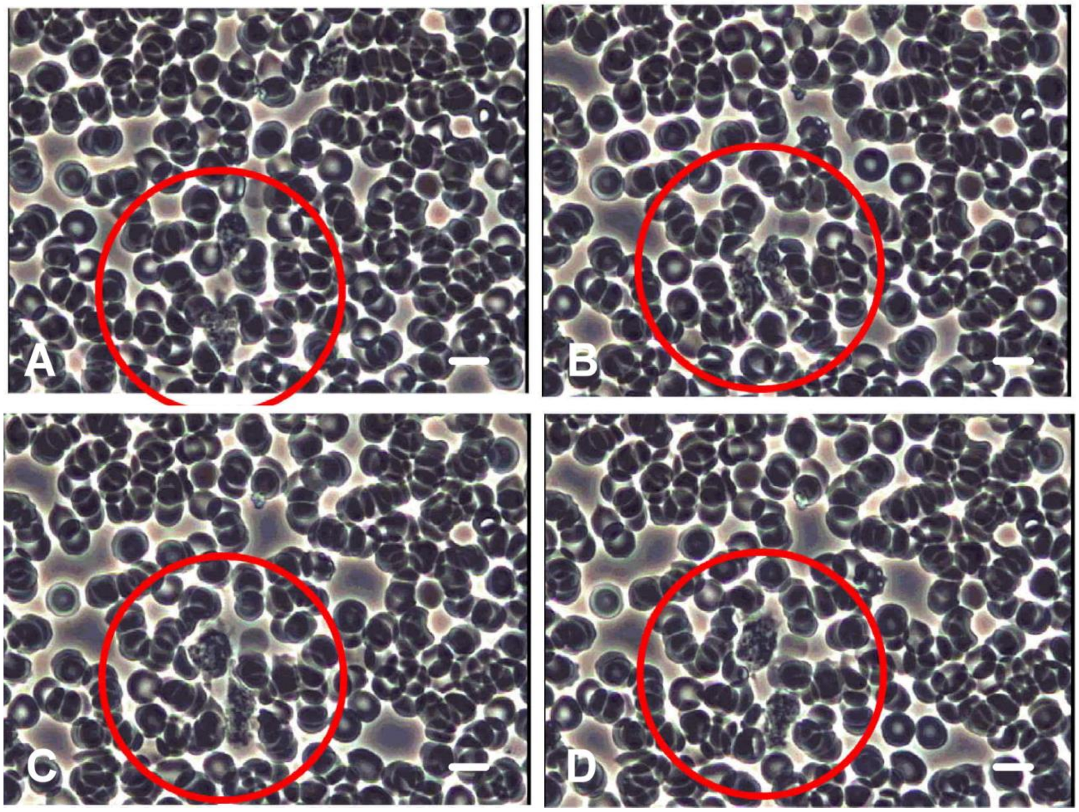


**Figure S3 The leukocytes in blood do not combine, versus Figure 2**

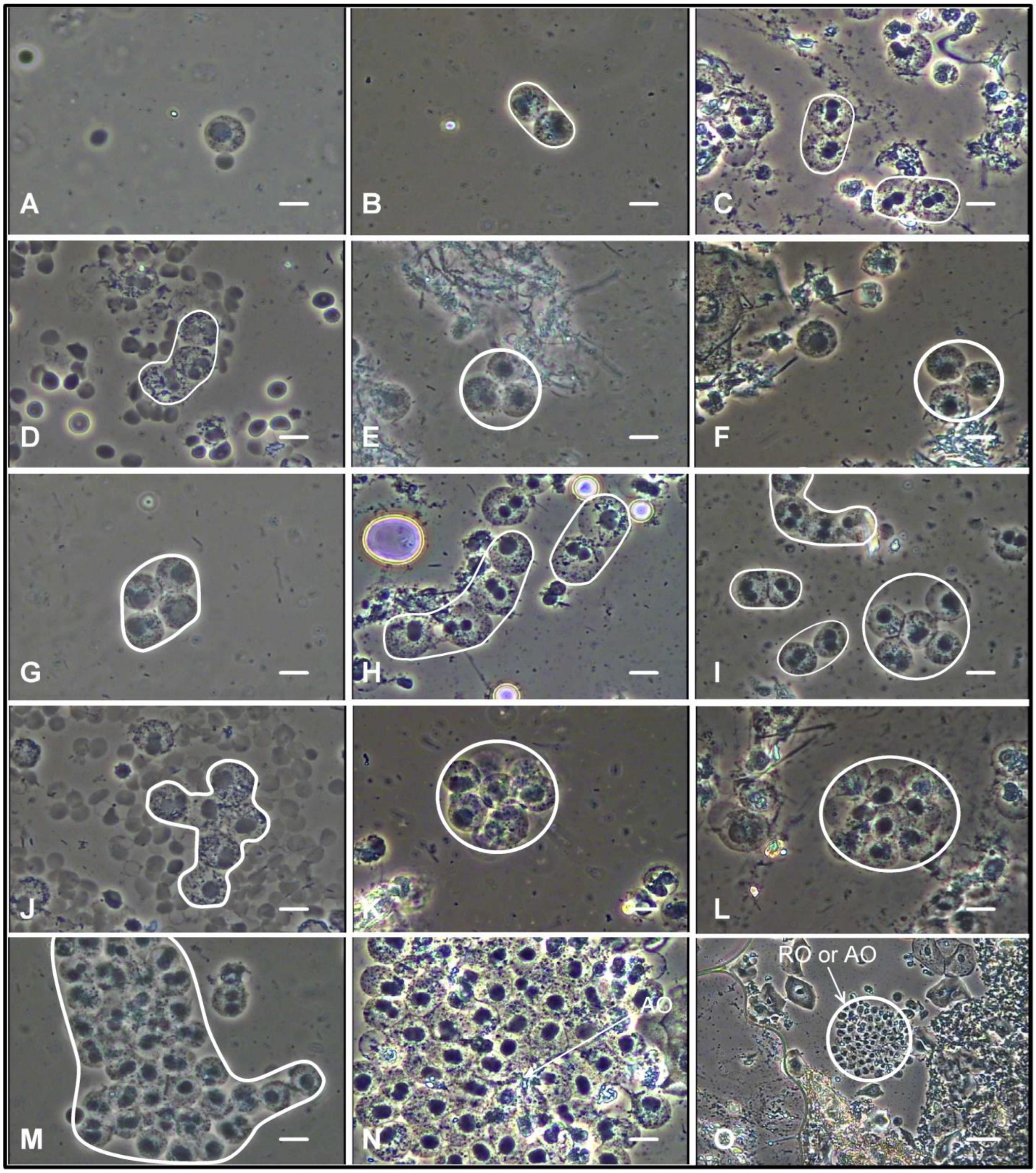


**Figure S4 The proportional size of the injury predicted by SLL of Figure 2**

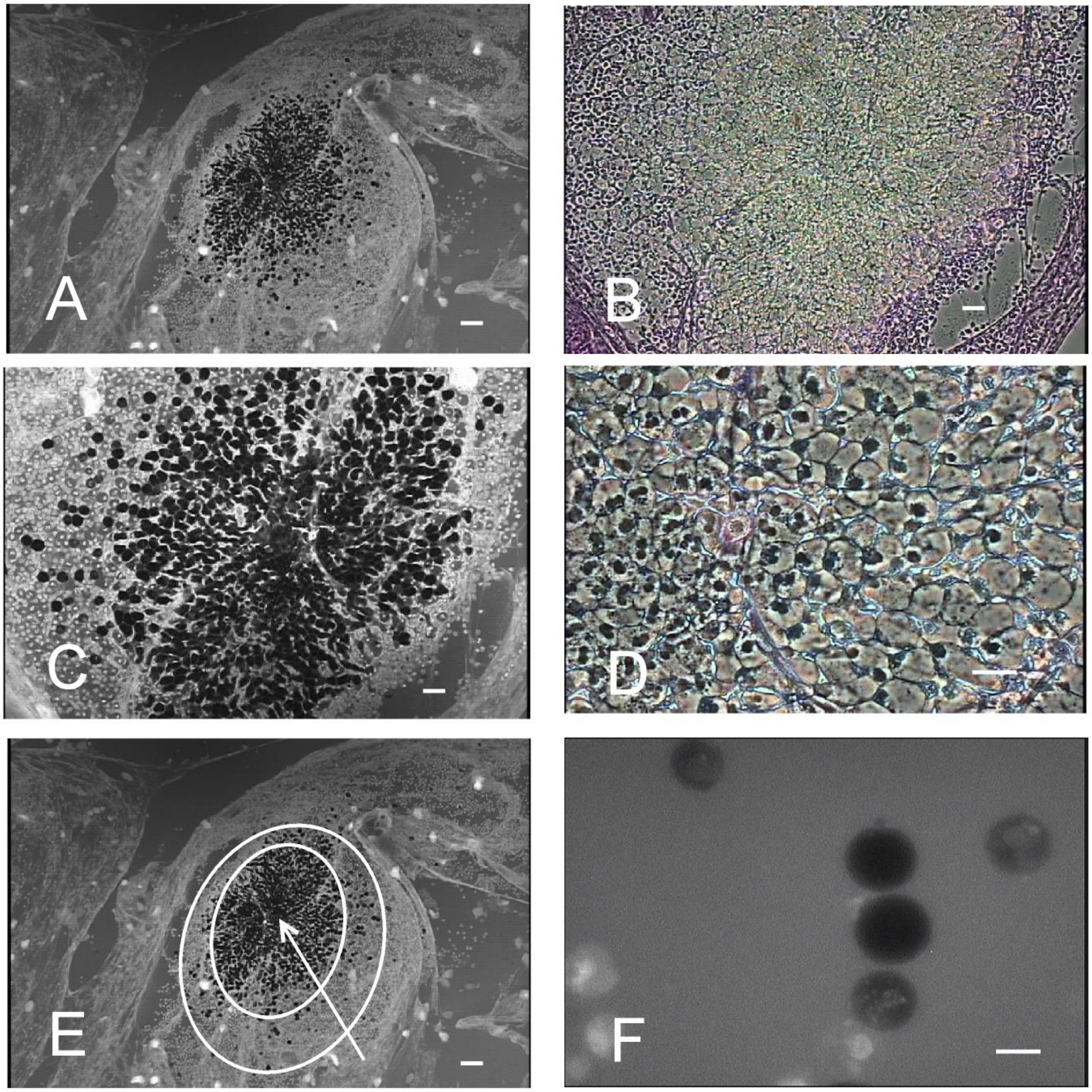


**Figure S5 MLC is SτC indicate space and time, as detail of Figure 5A, refer to Figure S6,**

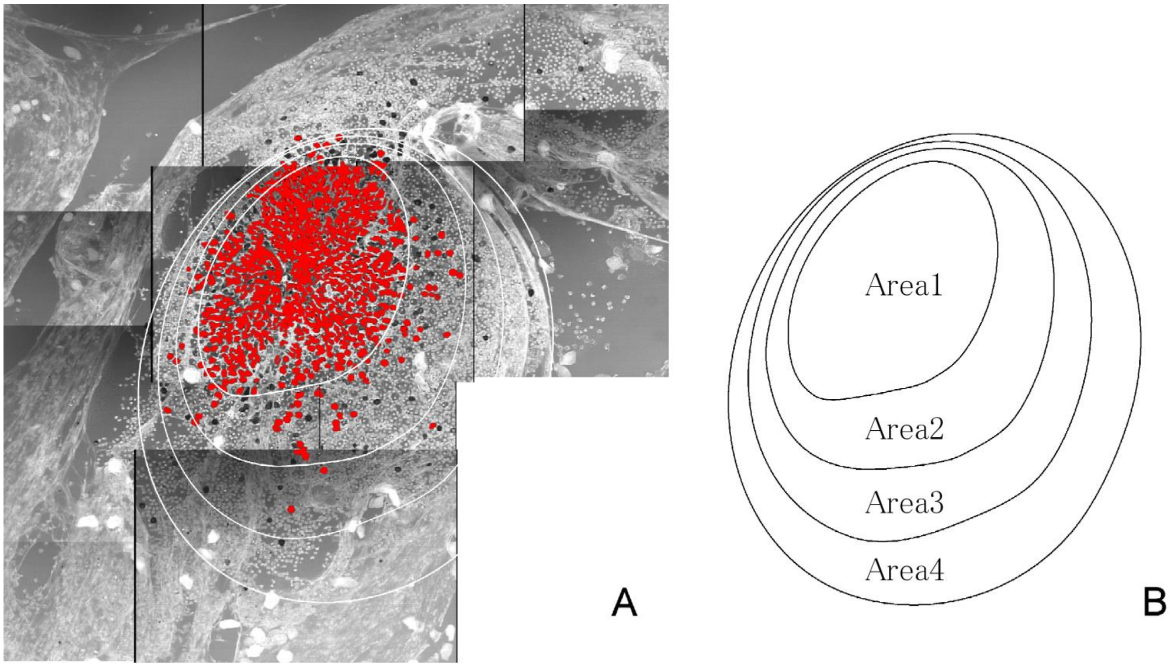


**Figure S6 For SOC and OOC, τ boundary and τ zone as Figure 5A LC**

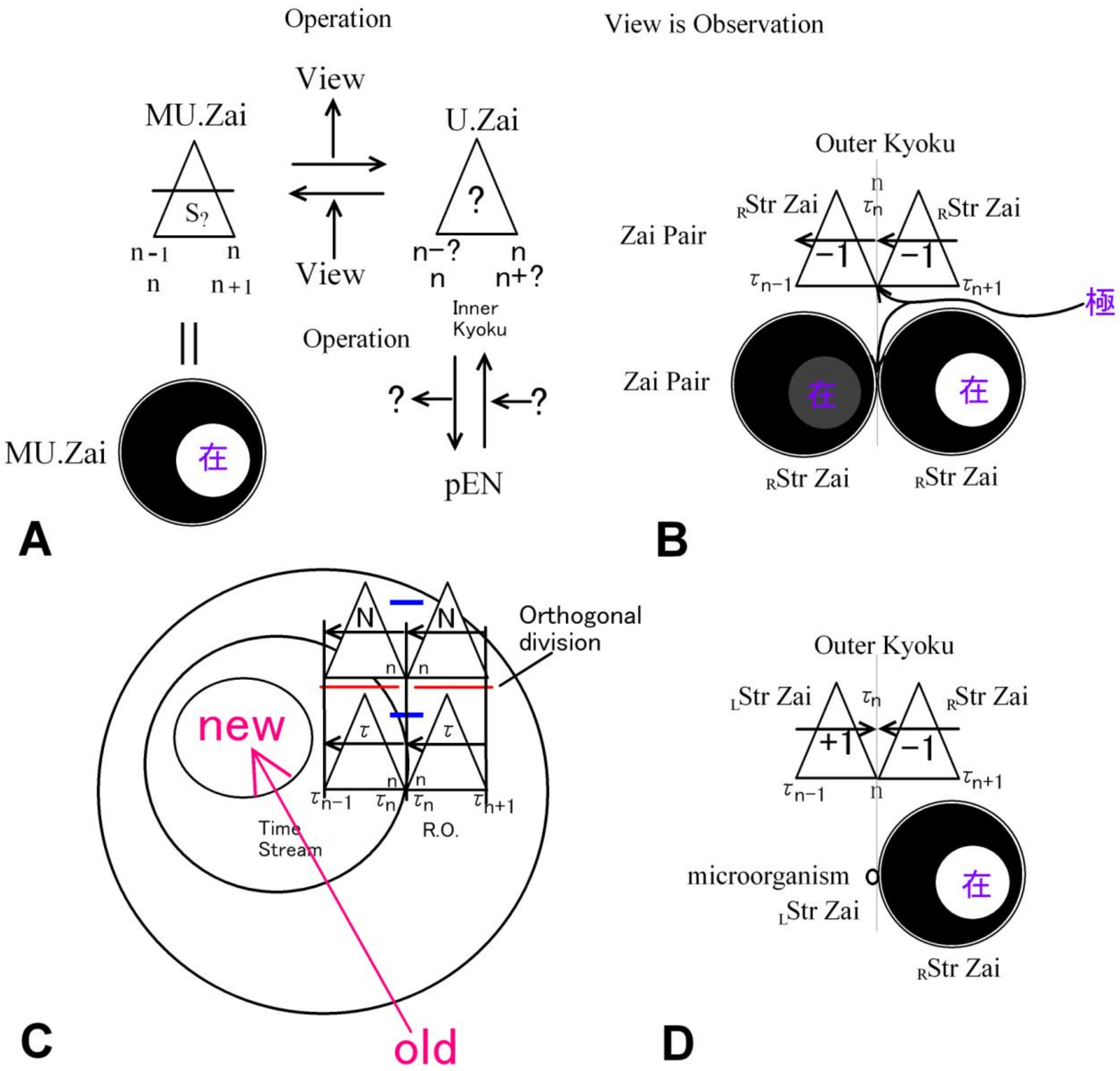


**Figure S7 The RDE as LC, for SOC and gsOOC in Figure 5A, S5, S6, and 7,**

SOC is formed as LC as SτC. The graph of a OOC by the SOC is indicated as red solid line (virtual line) as τSC. It is gsOOC. The SOC (Graph as the operation result) and gsOOC (Graph as the solution result) are calculation time zero. They are just P=NP.

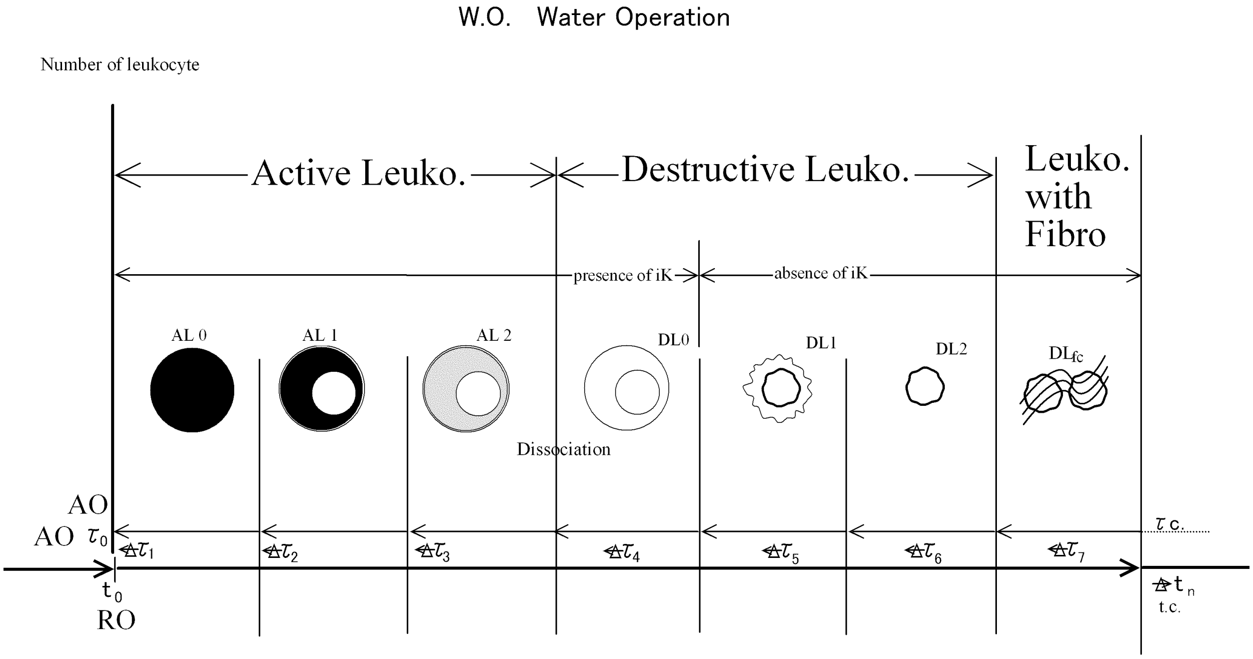


**Figure S8** water operation chart

iK is inner Kyoku, t.c is time continuum, τc. is τ continuum,

AL:Active Leukocyte, DL:Destructive Leukocyte, fc:Fibro is fibro tissue,

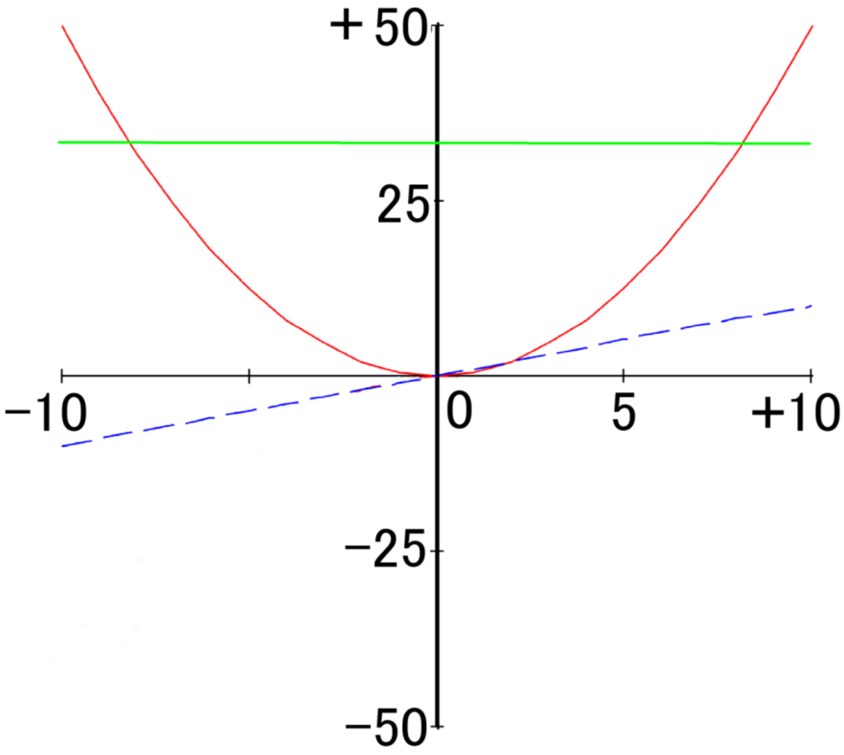


**Figure S9 Conventional equation of motion**

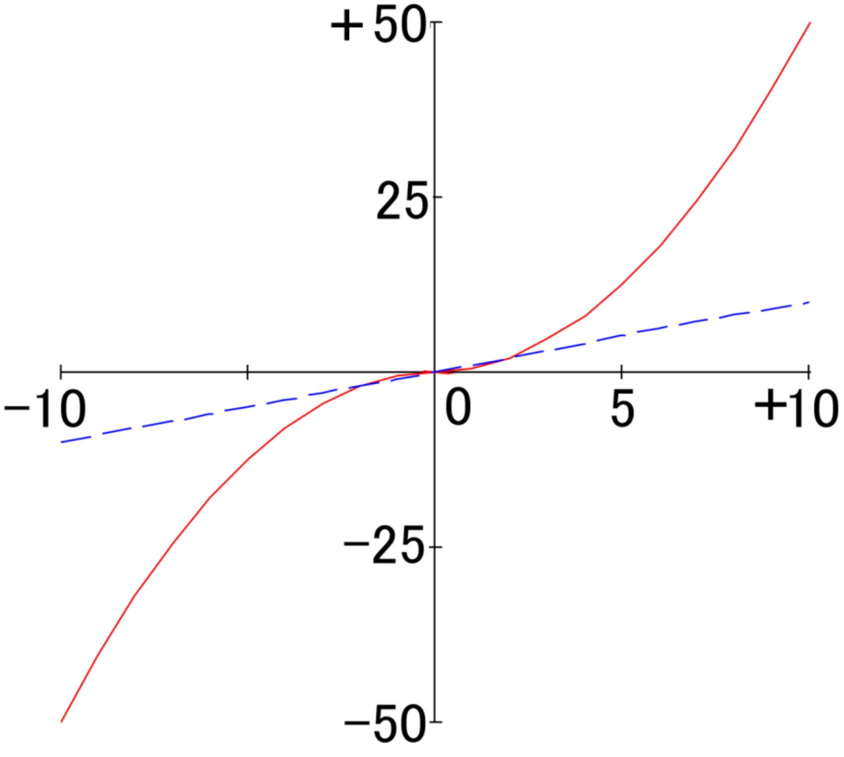


**Figure S10 Equation of motion with Str Operator (C3 coordinate system)**

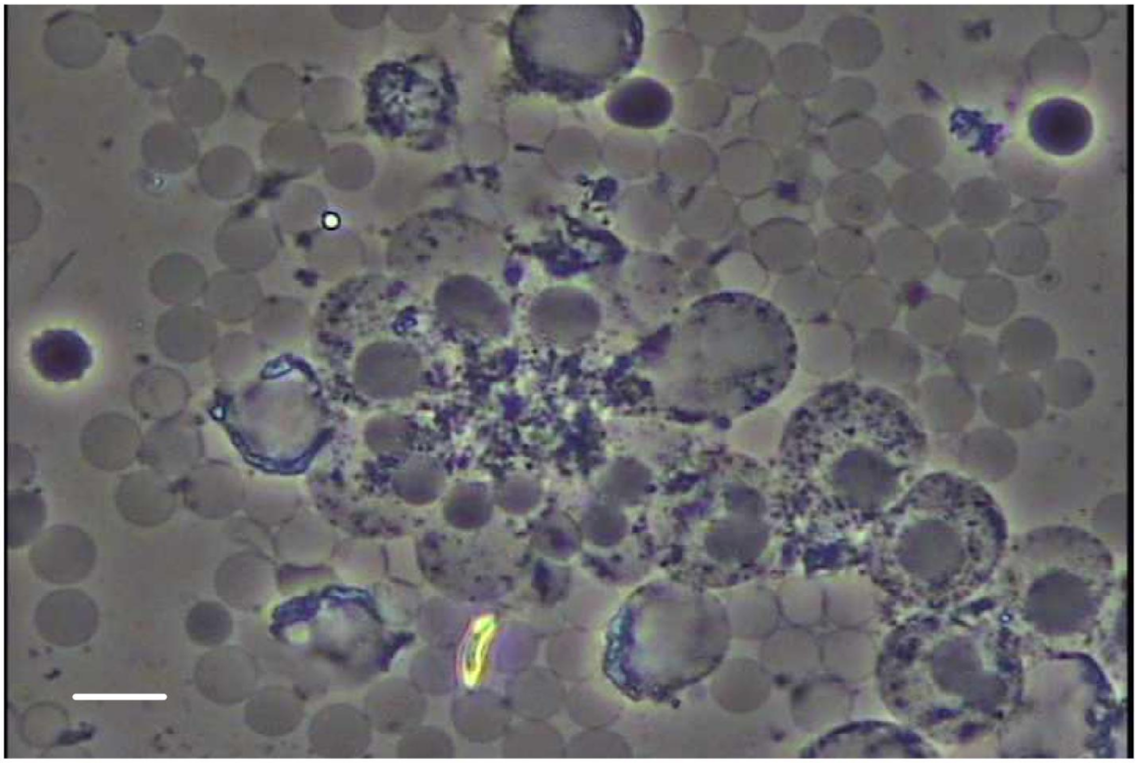

**Figure S11 Example of C4 Relative Differentiation and C4 RCS as Figure 7**

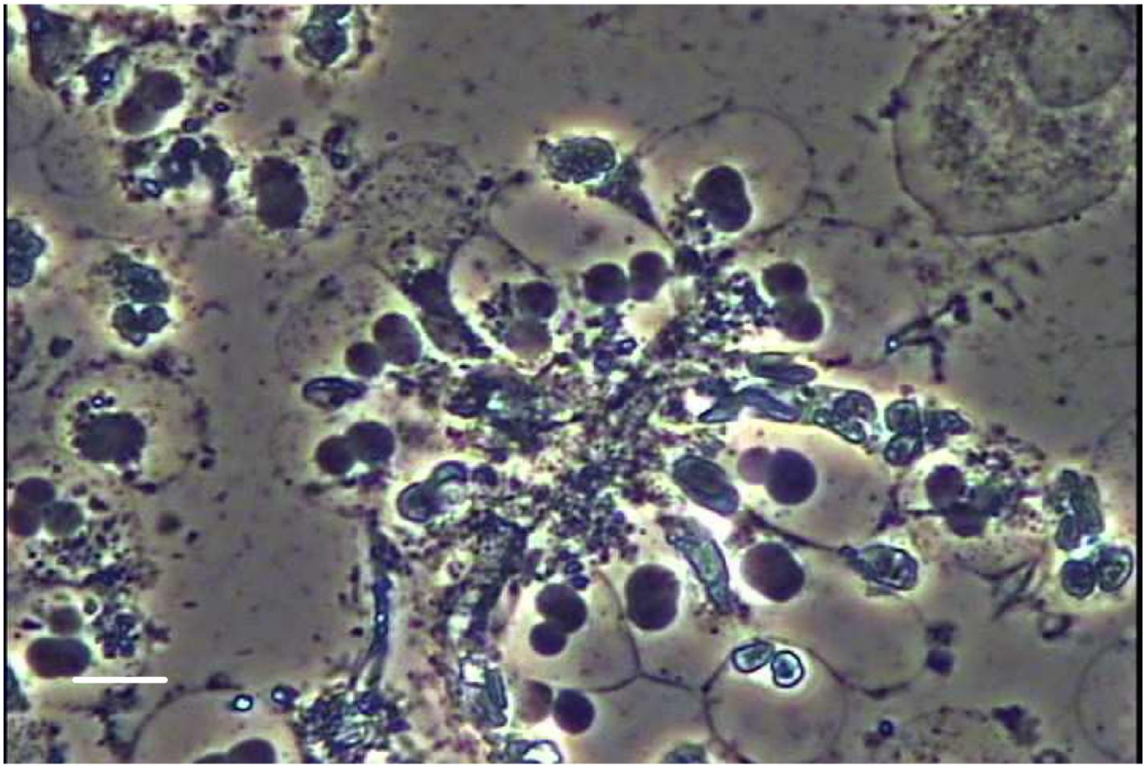

**Figure S12 Example of C4 Relative Differentiation and C4 RCS as Figure 7**

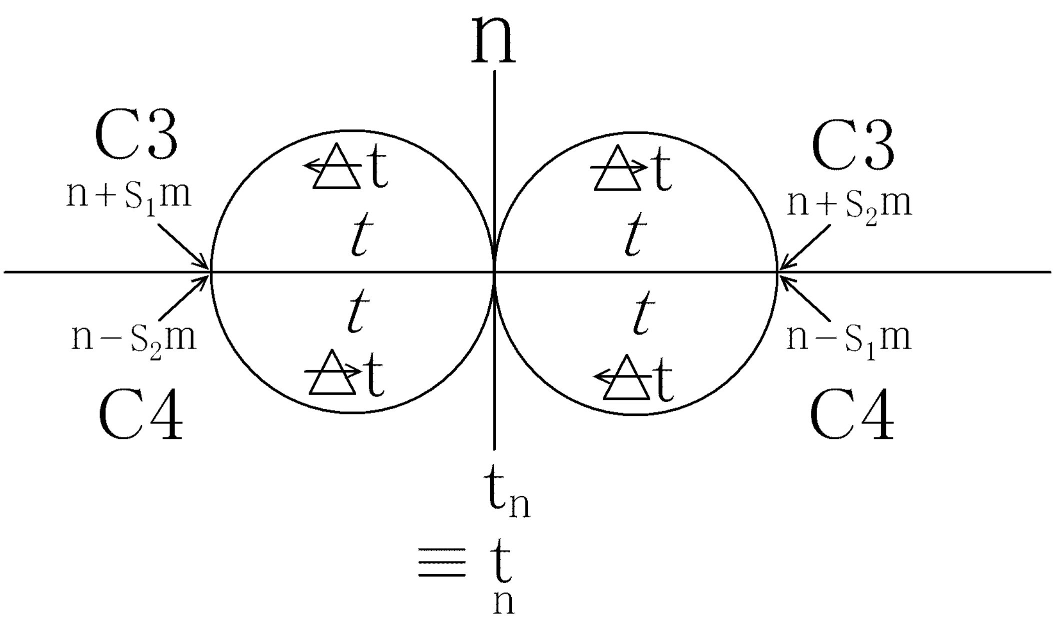


**Figure S13 physical time**

$\underset{n}{\mathrm{t}}$ and $\mathrm{t}_n$ is Kyoku time, t is pure number, $t$ is Zai time.
